# Personal clinical history predicts antibiotic resistance in urinary tract infections

**DOI:** 10.1101/384842

**Authors:** Idan Yelin, Olga Snitser, Gal Novich, Rachel Katz, Ofir Tal, Miriam Parizade, Gabriel Chodick, Gideon Koren, Varda Shalev, Roy Kishony

**Author notes:** Correspondence should be addressed to R. Kishony.

## Abstract

The prevalence of antibiotic resistance in urinary tract infections (UTIs) often renders the prescribed antimicrobial treatment ineffective, highlighting the need for personalized prediction of resistance at time of care. Here, crossing a 10-year longitudinal dataset of over 700,000 community-acquired UTIs with over 6,000,000 personally-linked records of antibiotic purchases, we show that the resistance profile of infections can be predicted based on patient-specific demographics and clinical history. Age, gender, and retirement home residence had strong, yet differential and even non-monotonic, associations with resistance to different antibiotics. Resistance profiles were also associated with the patient’s records of past urine samples and antibiotic usage, with these associations persisting for months and even longer than a year. Drug usage selected specifically for its own cognate resistance, which led indirectly, through genetic linkage, also to resistance to other, even mechanistically unrelated, drugs. Applying machine learning models, these association patterns allowed good personalized predictions of resistance, which could inform and better optimize empirical prescription of antibiotics.

---

The resistance of bacterial pathogens to commonly used antibiotics is a growing public health concern, threatening the efficacy of life-saving antibiotic drugs ^1,2^. Antibiotic use and misuse can benefit resistant strains, exacerbating the problem over time ^3–5^. At the single patient level, the efficacy of antimicrobial treatment is critically dependent on correctly matching antibiotic choice to the specific susceptibilities of the pathogen ^6–8^. Ideally, correct prescription should be based on direct measurement of the antibiotic susceptibilities of the infecting pathogen. In practice, though, to save time and resources, drugs are often prescribed empirically in absence of culture susceptibility measurements, risking incorrect and ineffective treatment.

This problem is of particular importance in Urinary Tract Infections (UTIs) which are often treated empirically despite substantial levels of resistance. UTIs are among the most common bacterial infections, with over 150 million annual cases globally ^9^. One of three women will have at least one symptomatic UTI by age 24, and more than one-half will be affected during their lifetime ^10^ Treatment of these infections accounts for about 8% of non-hospital usage of antibiotics, often as part of empirical prescription ^11,12^. The common etiological agents of UTIs are diverse, including *Escherichia coli, Klebsiella pneumoniae* and *Proteus mirabilis,* as well as gram-positive bacteria such as *Enterococcus faecalis* ^13–18^ These pathogens are often resistant to several antibiotics, with resistance rates of infections exceeding 20% for commonly used drugs ^14,17,19^, emphasizing the challenge of choosing the most appropriate antibiotic treatment for each patient ^20^.

The risk of an infection being resistant to different antibiotics can be correlated with patient demographics and clinical and epidemiological factors. Known demographic factors associated with resistance include older age ^21^, gender ^22^, ethnicity ^23–26^ and residence in a retirement home ^22^. Known clinical and epidemiological factors associated with resistance include presence of a urinary catheter ^18,22,27^, immunodeficiency ^22^, diabetes ^22^ and travel to developing countries ^25^ Notably, most of these associations were identified based on small patient cohorts, typically with high frequencies of antibiotic resistant infections, such as retirement homes, rehabilitation centers, or hospitals.

Beyond the patient’s demographics and current clinical factors, antibiotic resistance can also be associated with the patient’s past clinical history, including recurrent UTIs and hospitalizations as well as past resistance profiles and antibiotic prescriptions. Risk of resistance to specific drugs have been shown to increase for patients with recurrent UTIs ^22,26,28^ and past hospitalizations ^22,29^ Focusing on hospital settings, studies have further shown that past resistance profiles can be used to predict resistance in future infections ^30^ Availability of antibiotic purchase data reveals patterns of antibiotic use and misuse ^31^ and shows that risk of resistance increases with short-term prior use of antibiotics ^5,21,22,29,32–34^. At the population level, a recent large-scale study revealed correlations between risk of trimethoprim/sulfa resistance and the volume of past purchases of the same drug (cognate) as well as of other drugs of different pharmaceutical classes (noncognate) ^17^ Such associations of usage of a given antibiotics with future resistance to other antibiotics can appear indirectly through genetic linkage among resistance mechanisms, but mathematically resolving direct and indirect selection for resistance has been challenging. Negative associations, where drug use is anti-correlated with resistance, have also been observed, but it has been difficult to discern the direction of causality ^17,35^. Finally, the time extent of these positive and negative associations of resistance with prior antibiotic usage and prior resistant samples is not well resolved, and it is also unclear whether and how these associations vary across resistances to different antibiotics.

Here, we present a systematic big-data analysis of a large population of UTI patients to unravel predictive features of antibiotic resistance. We analyze a patient-level longitudinal dataset of community and retirement-home acquired UTI cultures collected by Maccabi Healthcare Services (MHS), Israel’s second largest Health Maintenance Organizations, serving a diverse population of ~2 million patients. We first analyze correlations between demographic factors and antibiotic resistance. Then, comparing resistance data of multiple infections from the same patient, we unravel long-term “memory” of resistance over time. We also combine these culture records with patient-linked records of antibiotic use to quantify the extent and time of direct and indirect impacts of antibiotic use on resistance at the single-patient level. Finally, we develop machine learning models combining these demographic and historical factors for personalized predictions of resistance.

## Results

We retrieved data of all positive urine cultures of MHS patients for the ten-year period between 01-July-2007 and 30-June-2017, as well as patient demographics and record of antibiotic purchases for these patients (Methods). Among all ~2 million MHS patients, there were 711,099 recorded positive urine samples from 315,047 patients total. For each positive sample, one or more bacterial species were isolated and characterized (total of 736,793). The dataset included the species identification of these isolates as well as VITEK2 antibiotic resistances reinterpreted in accordance with CLSI guidelines (Sensitive, Intermediate, and Resistant). For each UTI sample, we also defined “sample resistance” as the maximal resistance for each antibiotic across all isolates from the same sample (96.4% of samples were identified as single species and their resistance profile is simply defined as the resistance profile of their single isolates). In our analysis, we focus on the 16 most frequently measured antibiotic resistances (Table 1 and Supplementary Fig. 1). All of MHS’s country-wide clinical tests are performed centrally (Methods), allowing reliable comparison across patients and time.

**Table 1:** List of antibiotic resistances analyzed in the study

| **Antibiotics** | **Class** |
| --- | --- |
| Trimethoprim/Sulfa | DHFR inhibitor |
| Ciprofloxacin | fluoroquinolones |
| Ofloxacin |  |
| Nitrofurantoin | nitrofuran |
| Gentamicin | aminoglycosides |
| Amikacin |  |
| Meropenem | carbapenems |
| Ertapenem |  |
| Ampicillin | penicillins<br>(+/- β-lactamase<br>inhibitors) |
| Amoxicillin/CA |  |
| Piperacillin/Tazobactam |  |
| Ceftriaxone | cephalosporins |
| Ceftazidime |  |
| Cefuroxime – Axetil |  |
| Cephalexin |  |
| Cephalothin |  |

At the population level, resistance frequencies of different pathogens to different antibiotics varied mildly across time. Three species, *E. coli, K. pneumoniae* and *P. mirabilis,* account for 85% of isolates (70%, 10%, 5%, respectively; Fig. 1a). These pathogens varied in their resistance profiles (Fig. 1b). Notably, for almost any of the 16 considered antibiotics, the chance of resistant infection is significant, indicating that antibiotic treatment efficacy could often be undermined. Notable exceptions are amikacin, meropenem and ertapenem, for which resistance frequencies were negligible (<1% of UTIs); yet these are last-line antibiotics of restricted use. These population-level frequencies of resistance were fairly static over time and only mild changes were observed in certain antibiotics and specific species (Fig. 1c and Supplementary Fig. 2). The diversity of pathogens and resistance patterns underscores the widely accepted notion that antibiotic prescriptions must be tailored based on the resistance profile of the infection, motivating the development of methods to better predict resistance ^20^

**Figure 1:**
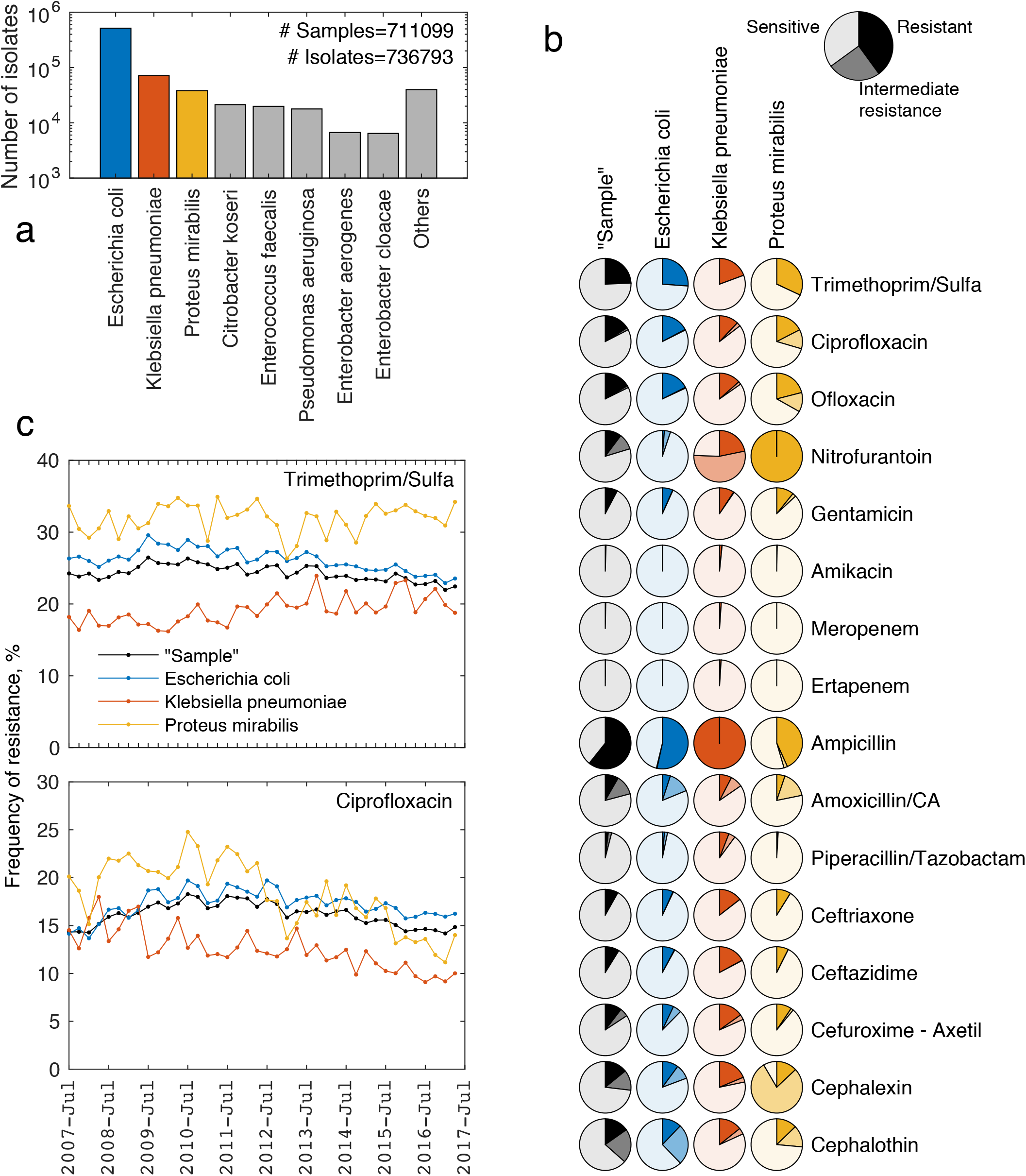
Frequency of bacterial species and antibiotic resistance in urinary tract infections. (**a**) Species abundance across the entire UTI dataset (July 2007-June 2017, 711099 samples, 736793 tested isolates). (**b**) The frequency of resistance and intermediate resistance to the 16 most frequently measured antibiotic drugs for the three most common bacterial species and for the urine sample as a whole (dark to light shades represent Resistant, Intermediate and Sensitive, respectively; Sample resistance is defined, for each urine sample as the highest resistance across all its isolates). (**c**) Frequencies of resistance for each of the three common species (colored lines) and the sample resistance (black lines) over the 10 year sampling time, for two representative antibiotics: trimethoprim/sulfa (top) and ciprofloxacin (bottom; See Supplementary Fig. 2 for all antibiotics.). Data points represent quarterly averages.

### The effect of demographics on antibiotic resistance

Considering the distribution of demographic factors across the UTI dataset, we find gender-specific age distributions (Fig. 2a). UTIs were much more common for females than males (~88% females, consistent with previous literature ^9,23^) and had qualitatively different age distributions. For females, UTIs appeared primarily in three distinct demographic groups: young children; reproductive-age women, including pregnant women; and the elderly, including retirement-home patients (Fig. 2a, top). In contrast, for males, there were only two peaks: an early-life peak of UTIs, dominated by infants, and a late peak of events for the elderly (Fig. 2a, bottom). Considering geography, the dataset spanned multiple cities which varied in their distribution of demographic factors (Supplementary Fig. 3). In total, this set of community-acquired UTIs showed diversity of demographic factors, allowing analysis of their impacts on resistance.

**Figure 2:**
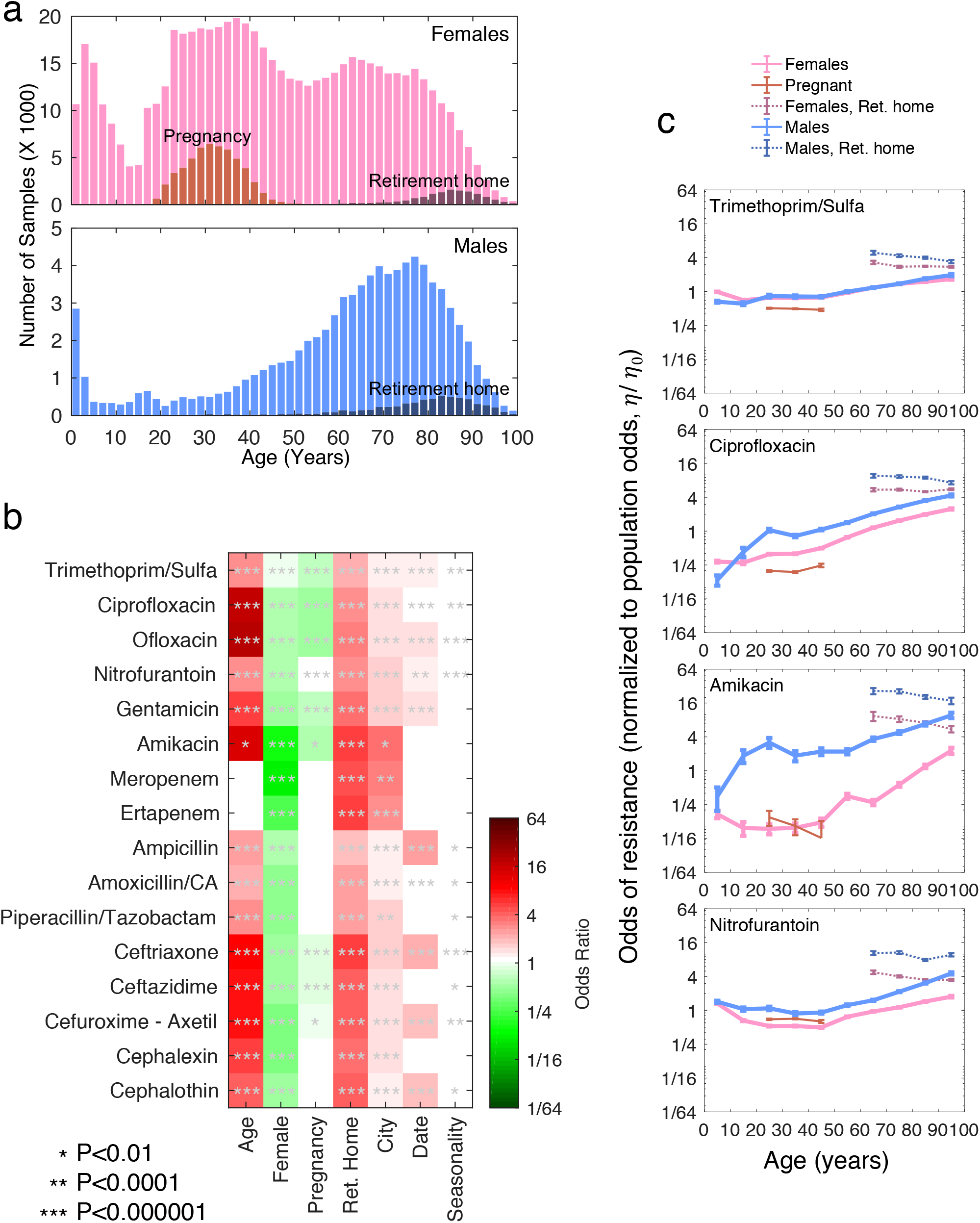
Strong and antibiotic-specific associations of resistance with demographic factors. (**a**) Distribution of positive urine cultures across major demographic factors: Age, Gender (top: Females, bottom: Males), Pregnancy (red) and Retirement home residence (black). (**b**) Adjusted odds ratios of resistance for each demographic variable (Methods: Logistic regression “Demographics”; for all adjusted and unadjusted regression coefficients see Supplementary Table 1). Asterisks indicate statistical significance and insignificant coefficients are shown as blank (white shade, P>0.01). (**c**) Odds of resistance as a function of age showing qualitatively distinct patterns for four representative antibiotics (Odds η are normalized to the average odds η_0_ across the population) UTI samples are separated into five non-overlapping categories: men not residing in retirement homes (blue), men residing in retirement homes (dotted blue), women not pregnant and not residing in retirement homes (magenta), women in retirement home (magenta dotted), and pregnant women (red). See Supplementary Fig. 4 for all antibiotics.

Age, gender, pregnancy and residence in retirement home had strong, yet differential, association with resistances to the 16 antibiotics. For each antibiotic, we performed multivariate logistic regression for the odds of resistance (*η* = *P_Resistance_*/(*P_sensitive_* + *P_intermediate_*) as a function of age (quadratically), gender, retirement home residence, pregnancy, city, date of sampling (quadratically) and season of sampling (Methods: Logistic regression “Demographics” model). For all of the 16 antibiotics, risk of resistance strongly increased with age and with retirement-home residence and decreased for females and pregnancy (Fig. 2b; and Supplementary Table 1 for regression coefficients and 95% CI). The effect of age on the odds of resistance differed widely among the 16 measured antibiotics, ranging from odds ratios (OR) of 2 in trimethoprim/sulfa and amoxicillin to more than 16 in antibiotics such as ciprofloxacin and amikacin (Fig. 2b and Supplementary Table 1). For most antibiotics, females had lower odds of resistance, yet the odds ratios varied substantially among the different antibiotics (from 0R=0.94 CI: 0.92-0.96 for trimethoprim/sulfa to 0R=0.12 CI: 0.10-0.15 for meropenem). These lower odds of resistance for females were even lower with pregnancy (as much as 0R=0.46; 95% confidence interval, 0.44-0.49 for ciprofloxacin; Supplementary Table 1). While the city of residence, when considered alone (Methods: Logistic regression “City, unadjusted”), was correlated with risk of resistance, these associations stemmed from differences in demographics among cities; they were much reduced in the complete regression model (compare adjusted and unadjusted coefficients, Supplementary Table 1). The date of sample had some association with resistance to specific antibiotics, most notably ampicillin and ceftriaxone, while season had almost no effect on resistance for any of the drugs (Fig. 2b).

The dependence of resistance on age showed complex and antibiotic-specific patterns. To better understand how the demographic factors interact and what is the effect of age as a continuous variable, we calculated, for each of the 16 antibiotics the odds of resistance *η* of the urine samples across ten age groups for five categories: men not residing in retirement homes, men residing in retirement homes, women not pregnant and not residing in retirement homes, women in retirement homes, and pregnant women (Fig. 2c and Supplementary Fig. 4). For some antibiotics, the odds of an infection being resistant were non-monotonic with age, with an additional peak of higher resistance risk at infancy or young age (e.g., nitrofurantoin, piperacillin/tazobactam; Fig. 2c and Supplementary Fig. 4). The effect of age was not always independent of other variables. We identified interaction between gender and age leading to heterogeneous patterns for males and females (e.g. amikacin, meropenem) and even to opposite effects of gender on odds ratios in specific ages groups (e.g. ciprofloxacin; Fig. 2c and Supplementary Fig. 4). While, across all antibiotics, resistance was higher for residents of retirement homes, the correlation with age within this group was reversed: the odds of resistance for retirement home residence decreased with age (Fig. 2c and Supplementary Fig. 4). We concluded that among the different demographic factors associated with risk of resistance, age, gender and residence in retirement homes are the strongest, with resistances to different antibiotics differentially affected by these factors and by interactions among them.

### Correlation of resistance among same-patient urine samples

Moving from demographics to clinical history, we analyzed correlations of resistance across same-patient infections, revealing short- and long-term “memory” and a constant patient-specific tendency for resistance. For each antibiotic A, we analyzed all same-patient pairs of samples and calculated the odds ratio for resistance of the second sample given the resistance of the first sample (*OR_pairs_* = (*N*_*R*→*R*_/*N*_*R*→*S*_)/(*N*_*S*→*R*_/*N*_*S*→*S*_), where the *N*’s are number of same-patient sample pairs with the specified resistance phenotypes; for example, *N*_*R*→*S*_ is the number of sample pairs in which the first sample is resistant to antibiotic A and the second sensitive; Methods). Calculating *OR_pairs_* as a function of the time difference *t* = *t*_l_ – *t*_2_ between the two samples in each pair, we find that, for all antibiotics, these odds ratios are highest for short time differences between the samples, then decay with two distinct time scales and finally converge, at the longer time differences, to a non-zero asymptotic effect (the odds ratios are well fitted by the sum of a fast exponent, a slow exponent and a constant, *OR_pairs_* ≃ *C*_0_*e*^*t*/τ_0_^ + *C*_0_*e*^*t*/τ_m_^ + *C*_∞_; Fig. 3a,b and Supplementary Fig. 5). The shortest time scale τ_0_, which is about a week, stems from successive samples likely taken from the same infection. For all antibiotics, resistance of a prior sample within this short time increased the odds of resistance of the second sample by not less than two orders of magnitude (Fig 3a,b,d, red). The second time scale τ_*m*_ represents much longer “memory” of correlations among samples, lasting more than six months for most antibiotics and even longer than a year for some (Fig. 3c). These long-term associations among sample measurements are consistent with repeated infection with the same or similar strains. The effect of this memory component was much more consequential and long lasting than previously thought. The maximal odds ratios of this memory term (*C*_m_) ranged between 6 and 14 for different antibiotics and typically remained larger than 2 even for samples taken a year apart (Fig 3a,b and Supplementary Fig. 5). At much longer times, this memory decayed, and *OR_pairs_* converged to a constant, but interestingly it did not diminish, but rather converged to values larger than 1 (Fig. 3a,b,d, green), representing timeless patient specific tendencies for resistance. In sum, we found an antibiotic-specific long-term memory of resistance between infections that can last for months to a year, as well as timeless correlations among even more remote samples, which can greatly improve predictions of resistance.

**Figure 3:**
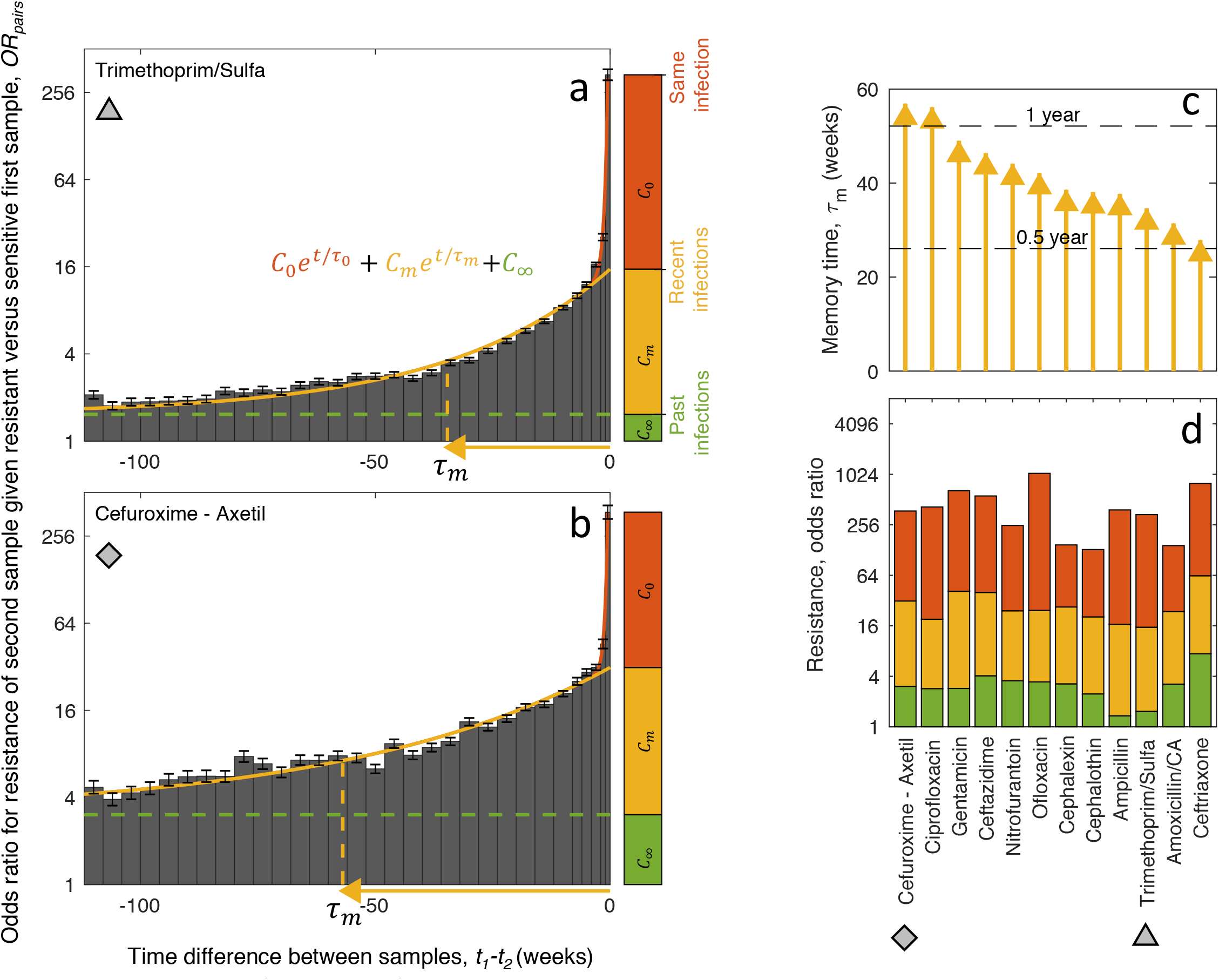
Long term “memory” of resistance across same-patient samples. (**a,b**) Odds ratio of resistance of a urine sample given resistant versus sensitive earlier sample from the same patient, as a function of the time difference between the two samples, for trimethoprim/sulfa and cefuroxime-axetil *OR_pairs_* = (*N*_*R*→*R*_/*N*_*R*→*S*_) / (*N*_*S*→*R*_/*N*_*S*→*S*_), Methods; Supplementary Fig. 5 for all antibiotics). Odds ratios are well fitted with *OR_pairs_* = *C*_0_*e*^*t*/τ_0_^ + *C*_0_*e*^*t*/τ_m_^ + *C*_∞_, representing short and long memory time scales and a timeless constant (red, yellow and green respectively). The magnitudes of these terms are shown as stacked bars on the right and the longer half life (τ_*m*_) is indicated across the time axis (yellow arrow). (**c**) Time scale of the long-term memory of resistance τ_*m*_ for different antibiotics. (**d**) The magnitude of short, long and timeless memory for different antibiotics (red, yellow, green bars).

### Association of resistance with past antibiotic purchase

Next, we crossed the infection dataset with patient-linked antibiotic purchase data. For each patient with recorded UTI samples, we retrieved all records of antibiotic purchase made during the twenty year period from 1-Jan-1998 to 30-Jun-2017. For analysis, we used 34 drug groups (Methods; Supplementary Table 2). For each UTI sample, we counted the number of purchases made by the same patient of each of the 34 drugs at distinct time intervals prior to the sample (Methods). Then, we applied multivariate logistic regression to correlate resistance to each of the 16 antibiotics with these drug purchase counts (Methods: Logistic regression “Purchase history”; Fig. 4a and Supplementary Fig. 6a).

**Figure 4:**
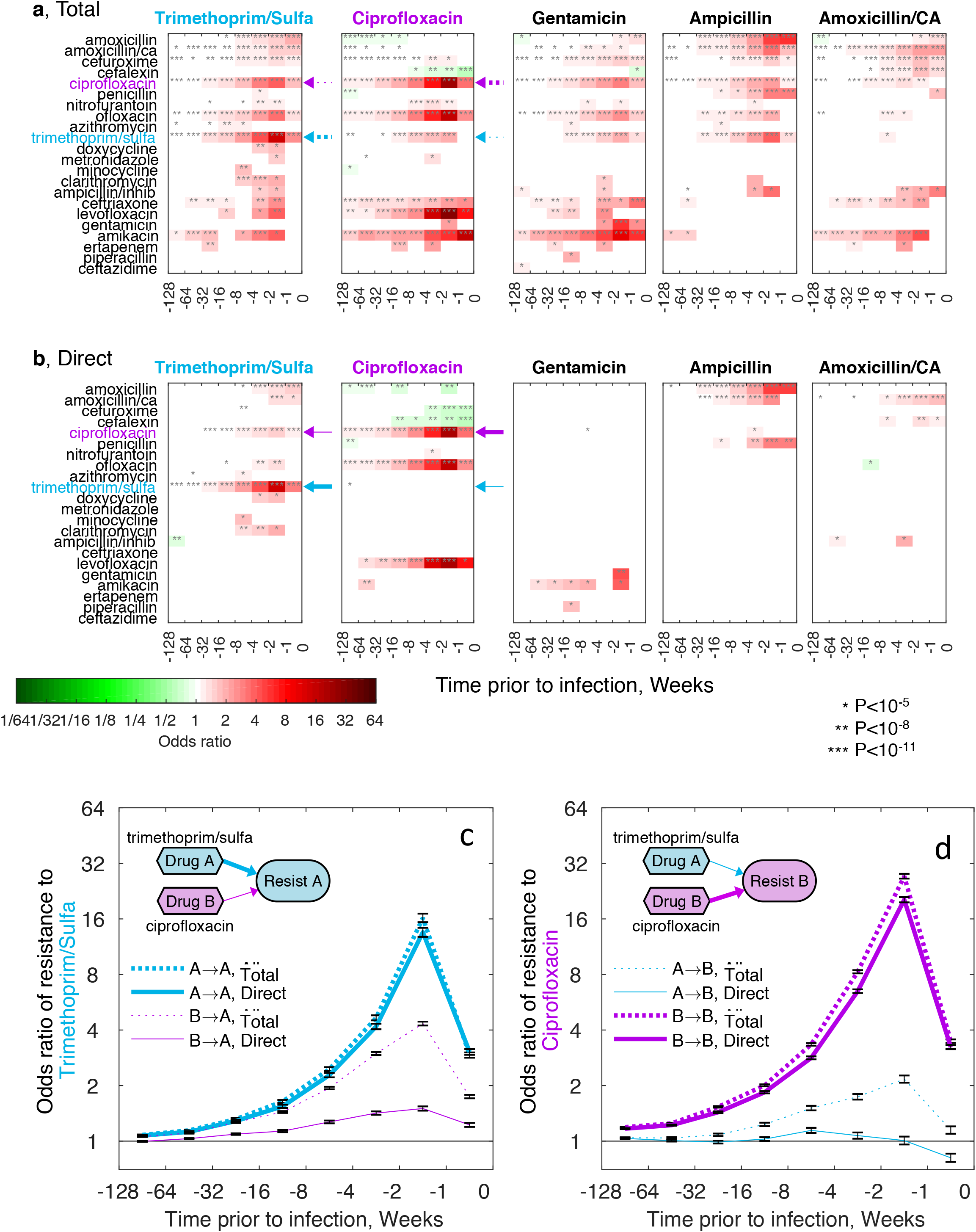
Past purchase of a given antibiotic has long-term direct association with its cognate resistance, which leads, through inherent linkage among resistances, also to indirect associations with other resistances. (**a**) Multivariate logistic regression models for association of resistance to 5 representative antibiotics (for all antibiotics see Supplementary Fig. 6a) with past purchases of different drugs at different time intervals prior to the time of sample (Methods: Logistic regression “Purchase history”; Values represent odds ratios for a single purchase of a specific drug at a specific time interval). Resistance shows long term association with past purchase of its matching antibiotic (cognate, see examples indicated by thick dashed arrows for trimethoprim/sulfa and ciprofloxacin) as well as with many other antibiotics (noncognate, see examples indicated by thin dashed arrows). (**b**) Logistic regression model as in (a) but, for each focal antibiotic A, adjusting for resistance of each urine sample to antibiotics different than A (Methods: Logistic regression “Purchase history adjusted for cross-resistance”; for all antibiotics see Supplementary Fig. 6c). This adjustment disentangles indirect effect where resistance to drug A is associated with purchase of drug B due to inherent association of resistance to B with resistance to A. This adjusted model is seen to diminish or even completely remove noncognate drug-to-resistance associations while mostly leaving the cognate associations un-attenuated. (**c,d**) Association of trimethoprim/sulfa and ciprofloxacin with purchases of their cognate drugs (cyan and magenta, respectively). Compare difference between total (Methods: Logistic regression “Purchase history”, dashed lines) and direct effect (Methods: Logistic regression “Purchase history adjusted for cross-resistance”, solid lines) for cognate (thick lines) versus noncognate (thin lines) drugs.

We identified strong long-term patient-level associations of resistance with past purchase of both cognate and noncognate antibiotics. For 22 out of the 34 drugs, we identified strong associations of prior drug purchase with future resistance, often lasting for months and even more than a year (Fig. 4a and Supplementary Fig. 6a). For example, the associations between purchase of ciprofloxacin and its cognate resistance having odds ratios of 1.5 after half a year and remaining as large as 1.2 even two years past purchase (Fig. 4a). Some weak negative associations were also identified (ciprofloxacin resistance was negatively correlated with past use of amoxicillin, Fig. 4a). Yet, the magnitude of these negative correlations decreased after adjusting for demographics, suggesting that they stemmed indirectly from correlations of purchases and resistance with demographics (Methods: Logistic regression, “Purchase history adjusted for demographics”; Supplementary Fig. 6b). Notably, drug purchases were associated not only with their expected cognate resistances. Indeed, use of both first-line antibiotics (such as ciprofloxacin, ofloxacin and levofloxacin) and last-line antibiotics (such as amikacin), increased the risk of a future resistance to a wide range of mechanistically diverse antibiotics. These abundant long-term positive associations between resistances and noncognate drugs did not stem from correlations of purchases and resistance with patient demographics; they remained strong even when adjusting for demographics (Supplementary Fig. 6b). Together, these results support strong and long-lasting associations of antibiotic resistance with past use of both cognate and noncognate antibiotics.

Exposing direct drug-to-resistance associations by disentangling linkage among resistances, showed that drug usage specifically selects for its cognate resistance at the single-patient level. Across the sample dataset, resistances to different antibiotics within class and even resistances to antibiotics of different classes were highly correlated (cross resistance; Supplementary Fig. 7). These inherent correlations among resistances suggest that observed associations between resistance to a given drug A and past purchase of a different drug B may arise indirectly through selection for resistance B and association between resistance to B and resistance to A. Mathematically discerning these direct from indirect effects is not possible when only a single resistance is considered ^17,36^ Yet, as our dataset contained measurements of multiple resistances for each sample, we were able to disentangle direct from indirect associations by adjusting the logistic regression for other measured resistances (Methods: Logistic regression “Purchase history adjusted for cross-resistance”). In this cross-resistance adjusted analysis of purchase-resistance associations, the noncognate associations between drug purchases and resistance much diminished and even disappeared while the associations between cognate drug-to-resistance pairs persisted (Fig. 4b and Supplementary Fig. 6c). For example, considering association between purchases of trimethoprim/sulfa and ciprofloxacin to their cognate resistances, we observed that the total (unadjusted) effect of cognate drugs remained when adjusting for cross-resistance (Fig 4c,d, thick solid vs. thick dashed lines), while the total effect of drugs on their noncognate resistance was much diminished when removing indirect effect (Fig 4c,d, thin solid vs. thin dashed lines). Our analysis therefore suggests both direct and indirect selection for resistance at the single-patient level lasting months and even a year following drug use.

### Predicting antibiotic resistance at the single-patient single-infection level

Finally, as resistance is strongly associated with demographics, sample history and purchase history, we wanted to determine the predictive power of these factors individually and when combined together. Models of Logistic Regression and Gradient Boosting Decision Trees (GBDT) were trained and tested on temporally separate periods: training period of 9 years from 1-July-2007 to 30-June-2016 and testing period of the following year, from 1-July-2016 to 30-June-2017 (for some antibiotics, training and testing periods were modified to avoid time periods during which resistance was not routinely measured, Supplementary Fig. 1, green/red horizontal bars). This temporal separation between training and testing data emulates forecasting resistance, as would be the case in real-life implementation of such a method. Area Under the Curve (AUC) of Receiver Operating Characteristic was used as a standard measure for predictive power ^37^.

Individually considering demographics, sample history and purchase history, we find that each of these sets of features had significant predictive power, yet their dominance varied across the different antibiotics (Fig. 5). For demographic parameters (Methods: Logistic regression “Demographics”), AUC ranged from barely 0.58 for ampicillin to values as high as 0.85 for amikacin, reflecting their weak and strong demographic dependencies, respectively (Fig. 2b). For sample history (Methods: Logistic regression “Sample history”), the predictive power was more consistent, with AUC varying between 0.64 for piperacillin/tazobactam to 0.83 for ertapenem. Purchase history (Methods: Logistic regression “Purchase history”) had barely any predictive power in drugs like meropenem and nitrofurantoin (AUC of 0.52 and 0.59, respectively), yet was higher for other drugs, especially for the fluoroquinolones (AUC of 0.78 for ofloxacin and 0.75 for ciprofloxacin). Comparing the relative strengths of these feature sets, we find that while they vary in dominance across the antibiotics, resistance to any given antibiotic was often well predicted by more than one set, suggesting that even higher predictability could be achieved by combining them.

**Figure 5:**
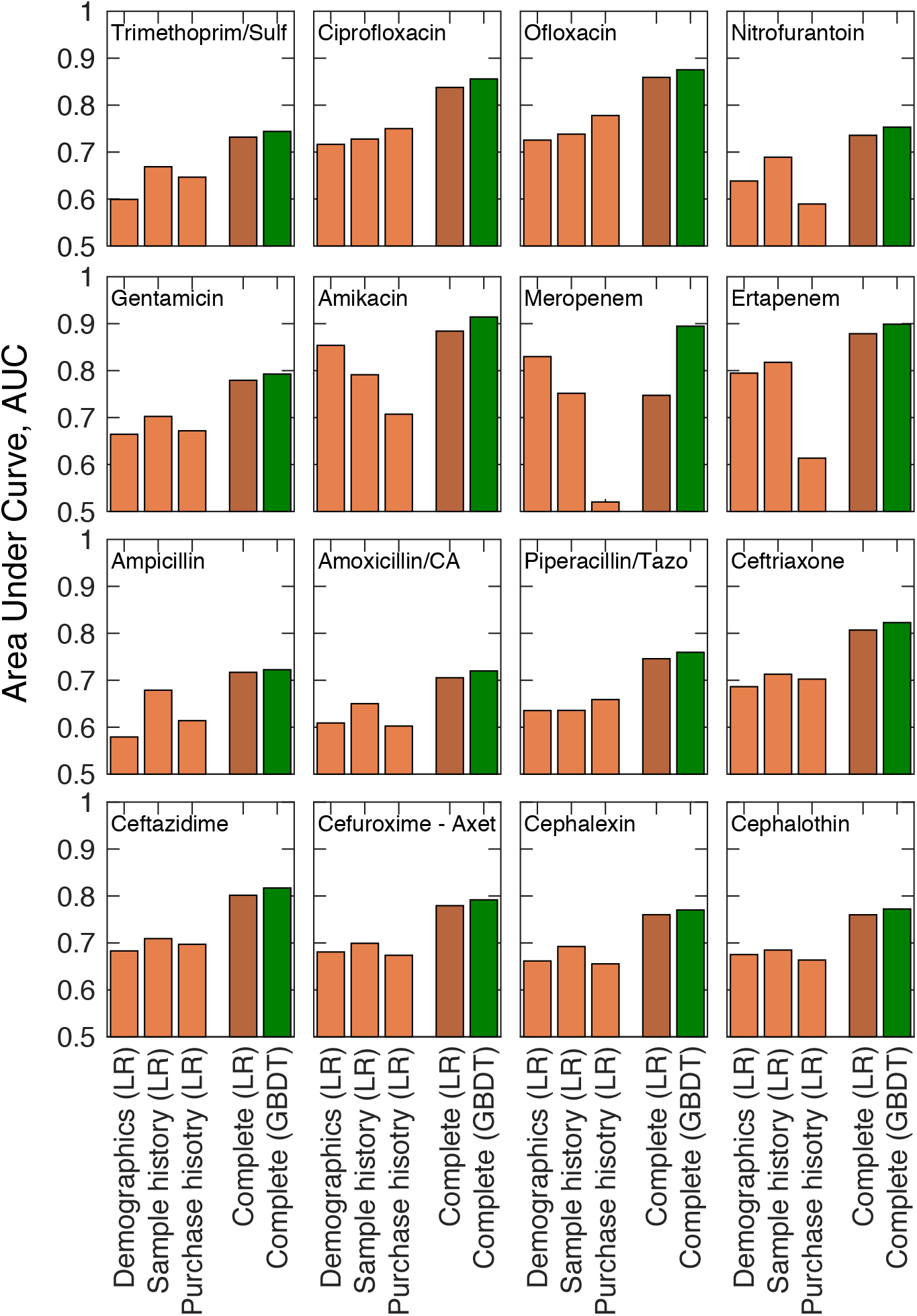
Prediction of resistance at the single-infection level based on patient demographics, urine sample history and drug purchase history. Area Under the Curve (AUC) of Receiver Operator Characteristic for prediction of resistance to each antibiotic using demographics, sample history and purchase history individually in a Logistic Regression model (LR, light orange), as well as when all of these feature sets are combined together in LR (dark orange) or in Gradient Boosting Decision Trees (GBDT, green). Models were trained on urine samples within the training date range (typically 9 years; see Supplementary Fig. 1) and tested on the urine samples collected during the year following the training period (green and red horizontal bars, Supplementary Fig. 1).

Combining these feature sets, using logistic regression or GBDT models, much increased predictability of resistance. Using all features together in a complete logistic regression model (Methods: Logistic regression “Complete”), provided good to excellent predictability to all antibiotics (AUC ranged from 0.7 for amoxicillin/CA to and beyond 0.85 for ertapenem, amikacin and ofloxacin; Fig. 5). Predictability of resistance across all antibiotics was increased even further by the complete GBDT models (Methods; Fig 5), most notably for meropenem, which was prone for overfitting (Supplementary Fig. 8). A closer look at the decision trees of the GBDT models identified the relative importance of the different features (Supplementary Fig. 9). While some features, like age and past resistance were important in all models, others like gender and time interval from last antibiotic purchase where more specific to each resistance. Together, these results demonstrate how machine learning models, trained on the combined dataset of demographics, sample history and drug purchase history, can provide high predictability of antibiotic resistance at the single-patient and single-infection levels.

## Discussion

Analyzing a large longitudinal medical dataset, we demonstrate the predictability of antibiotic resistance in UTIs based on demographics, antibiotic resistance profile of past UTIs and purchase history of antibiotic drugs. These culture-free predictions could guide appropriate antibiotic prescription. Considering demographics, we found that - age, gender, pregnancy, and residence in a retirement home were strongly associated with resistance, showing complex even non-monotonic patterns specific to each of the different antibiotics. Utilizing repeated same-patient cultures in our database, we identified and characterized a personal component of “memory” of resistance, lasting for many months and even over a year and further contributing to predictability of resistance. Long memory was also observed between resistance and past drug purchases. Resistance to a given drug had long-lasting associations not only with past usage of this same drug, but also with other, even mechanistically unrelated, drugs. Yet, adjusting for linkage among resistances exposed direct selection where drug use leads specifically to its own cognate resistance at the single patient level. Taken together, these results are consistent with drug use directly selecting, at the single-patient level, for strains resistant to it and thereby selecting indirectly, likely through genetic linkage, to resistance to other antibiotics. Once resistance strains occurs they persist in the body affecting the chance of resistant infections months ahead.

Some aspects of the data may complicate the interpretation of our results, especially wherever causality may be inferred. As purchase of a drug does not fully guarantee its concurrent use, later usage of a purchased drug may bias our results towards higher odds ratio for purchases made long before infection. Conversely, we can not exclude that some patients have used antibiotics they did not purchase through MHS, which will bias our results towards lower odds ratio for drug purchases. Also, although culture data is routine for suspected UTIs, sending urine for a culture test is not obligatory. As a result, we assume some UTIs would be empirically treated without any culture record, and there could be enrichment for culture tests following treatment failure. Such enrichment towards cultures following treatment failure can generate bias towards measurements of more resistant samples, resulting in overestimation of the total frequency of resistance, especially for first-line treatment. It can further contribute to the strong short-term association of drug purchases with resistance, especially for first-line antibiotics. The level of enrichment for urine culture tests following treatment failure could itself depend on demographics, which can bias correlations of demographics with resistance. While we cannot exclude these biases, our analysis firmly demonstrates that, with all of these potential biases, resistance of urine samples can be well predicted based on the patient’s specific demographics and clinical history.

The predictability of infection resistance profiles lays the basis for a future paradigm where clinicians will routinely consult machine learning algorithms for prescription of antibiotic treatment tailored by the patient records and their clinical history. While the key factors identified here can serve as the basis of such approach, we expect that the specific model, the exact coefficients and relative weights of predictors, must be tailored for each country or region. Indeed, these algorithms can also be dynamically and adaptively updated in real time as new data is acquired. We also expect that inclusion of additional patient specific factors, such as comorbidities, can further increase resistance predictability. In the longer term, these clinical-record based approaches could be integrated with genomics of the patient as well as of the pathogen ^38–244^ Implemented in the clinic, machine-learning guided personalized empirical prescription can reduce treatment failure as well as lower the overall use and misuse of antibiotics thereby assisting in the global effort of impeding the antibiotic resistance epidemic.

## Methods

### Data

Anonymized clinical records of urine culture tests (“culture reports”) and records of antibiotic purchases (“purchase reports”) were obtained from Maccabi Health Services (MHS) for the time period from July 2007 to June 2017. Random patient identifiers were used to link culture reports and antibiotic purchase reports.

#### Culture reports

Antibiotic resistance profiling of bacterial pathogens isolated from urine cultures was carried out centrally (in two locations until 2010, and in one central lab since). We retrieved 711,099 culture reports of positive samples from 315,047 patients total. Each report included: (1) Unique patient code; (2) Date of sample; (3) List of isolates cultured with species identification (typically one isolate per sample; 3.6% of samples had more than one isolate); (4) Resistance profile of the isolates from processed results of a VITEK 2 system given as Sensitive, Intermediate and Resistant for each drug tested. Not all antibiotics were measured for each sample, and only the 16 most commonly measured antibiotics were included for further analysis (*N_Resistances_* = 16, Supplementary Fig. 1, Table 1). Additionally, for each of the 16 included antibiotic resistances, we removed time intervals where only a small percentage of samples were measured to avoid sampling bias (Supplementary Fig. 1, gray). (5) Demographics: age, gender, pregnancy of the patient, as well as a code for the reporting physician which we used to identify patients residence in retirement homes. (6) Referring lab identifier, which was used to assign patients to a city of residence.

#### Antibiotic purchase reports

All drug purchases by prescription are routinely recorded in MHS databases. We identified and retrieved all purchases by patients with culture reports by converting internal MHS drug codes to ATC classifications of antibiotics (Supplementary Table 2). Each purchase record included: (1) Unique patient code to be linked to the code of the culture record; (2) Internal MHS product code, which was translated to an ATC drug code, (3) Date of purchase.

#### Choice of drugs for analysis

We focused on 34 groups of antibiotic compounds (*N_ATC_* = 34), including the 30 most purchased during the time of the study (groups 1-30, Supplementary Table 2) as well as less frequently purchased groups of compounds manually identified as cognate drugs of the 16 antibiotic resistances analysed (groups 31-33 in Supplementary Table 2).

### Feature definition

For each cultured sample, we define the following parameters used for the logistic regression and the gradient boosting decision trees:

#### Sample resistance profile

For each urine sample, we define *Y_k_* as 0 for sensitive and intermediate and 1 for resistant to antibiotic *k* (1 ≤ *k* ≤ *N_Resistances_*). If the sample had multiple isolates, *Y_k_* was assigned 1 if at least one isolate was resistant. Missing resistance measurements are defined as N/A, and for each antibiotic *k* only samples which have defined resistance to it are used when training or testing its Logistic Regression or Gradient Boosting Decision Trees (GBDT).

#### Demographics

*X^Gender^*: 0/1 for males/females; *X^Pregnancy^*: 0/1 indicating pregnancy; *X^RetHome^*: 0/1 indicating residence in retirement homes; *X^Age^*: patient age in years at time of UTI sampling; *X^Date^*: date of sample in days starting at 1-July-2007; *X^Season^*: the day of sample within the calendar year, where 1 is 1-Jan and 365 is 31-Dec; 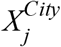: 0/1 indicating residence in city *j* (1 ≤ *j* ≤ *N_City_*, *N_City_* = 20, Supplementary Fig. 2 for City list; we use Tel-Aviv as a reference); for GBDT, nominal *X^City^* ∊ {1, ‥,20} was used instead.

#### Sample history

For a given sample, we consider all earlier samples taken from the same patient (if any). We bin the time difference between any such earlier sample and the current sample, *t* = *t_past sample_* – *t_Sample_* (*t* is negative designating past events), into one of 8 logarithmically spaced time bins (*i* = 1,2,…,*N_time bins_ N_time bins_* = 8 a bin *i* is defined by *t_i_* ≤ *t* < *t*_*t*–1_, where the boundaries of these time bins are *t*_0_ = 0, *t*_1_ = −1, *t*_2_ = −2, *t*_3_ = −4,…, *t*_8_ = −128 weeks). We then calculated 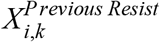 and 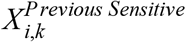 as the number of prior cultures within time bin *i*, with sample resistance *Y^k^* equal to 0 or 1 (Resistant or Sensitive), respectively.

#### Drug purchase history

For each urine sample, we consider all earlier drug purchases made by the same patient. We bin the time difference between the urine sample date and a given past purchase, *t* = *t_Purchase_* − *t_Sample_*, into 8 time bins as above. For each sample, we then calculate 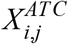 as the number of purchases of drugs of a given group *j* (1 ≤ *j* ≤ *N_ATC_*, Supplementary Table 2) made by the patient during time bin *i*. For the GBDT model, we also included as additional features the types *j* of the 3 latest purchases (nominal features) and their purchase-to-sample time difference (*t*, in days)

#### Cross-resistance

To resolve direct versus indirect associations of drug purchase and resistance, we adjusted the logistic regression of resistance to a given antibiotic *k* as a function of past drug purchases by the resistances to all other drugs *j* which are non-analogous to *k*. We define *A_k,j_* as a binary variable equals 0 and 1 for analogous versus non-analogous drug pairs, respectively. “Analogous” pairs are defined as antibiotics which have exceptionally high cross-resistance (*A_k,J_* = 0 for *Corr*(*Y_k_*, *Y_j_*) > *A_threshold_*; we use *A_threshold_* = 0 7 which corresponds to drug pairs of the same class; see pairs labeled with ‘x’ in Supplementary Fig. 7). We then add as features in the regression analysis of a given antibiotic *k* the resistance measurements *Y_j_* to all antibiotics *j* for which *A_k,j_* = 1. <u>Note</u>: These cross-resistance features provide information from the focal sample and were used only in the analysis of direct/indirect effect of purchases (Fig. 4 and Supplementary Fig. 6c). Importantly, they were not used when evaluating predictability of resistance (Fig. 5).

### Logistic regression

Logistic regression of resistance for each antibiotic was performed via the Matlab glmfit function. For each of the 16 antibiotics *k*, the probability of resistance *P_k_* was fit to the sample resistance *Y_k_* for all urine samples which had measurement of resistance to *k* either across the entire 10 year dataset (for Figs. 2,4), or across the “training period” (for the analysis of predictive power of Fig. 5; see Supplementary Fig. 1 for definition of the training period for each of the 16 antibiotics). The different logistic models included combinations of the following 10 terms:

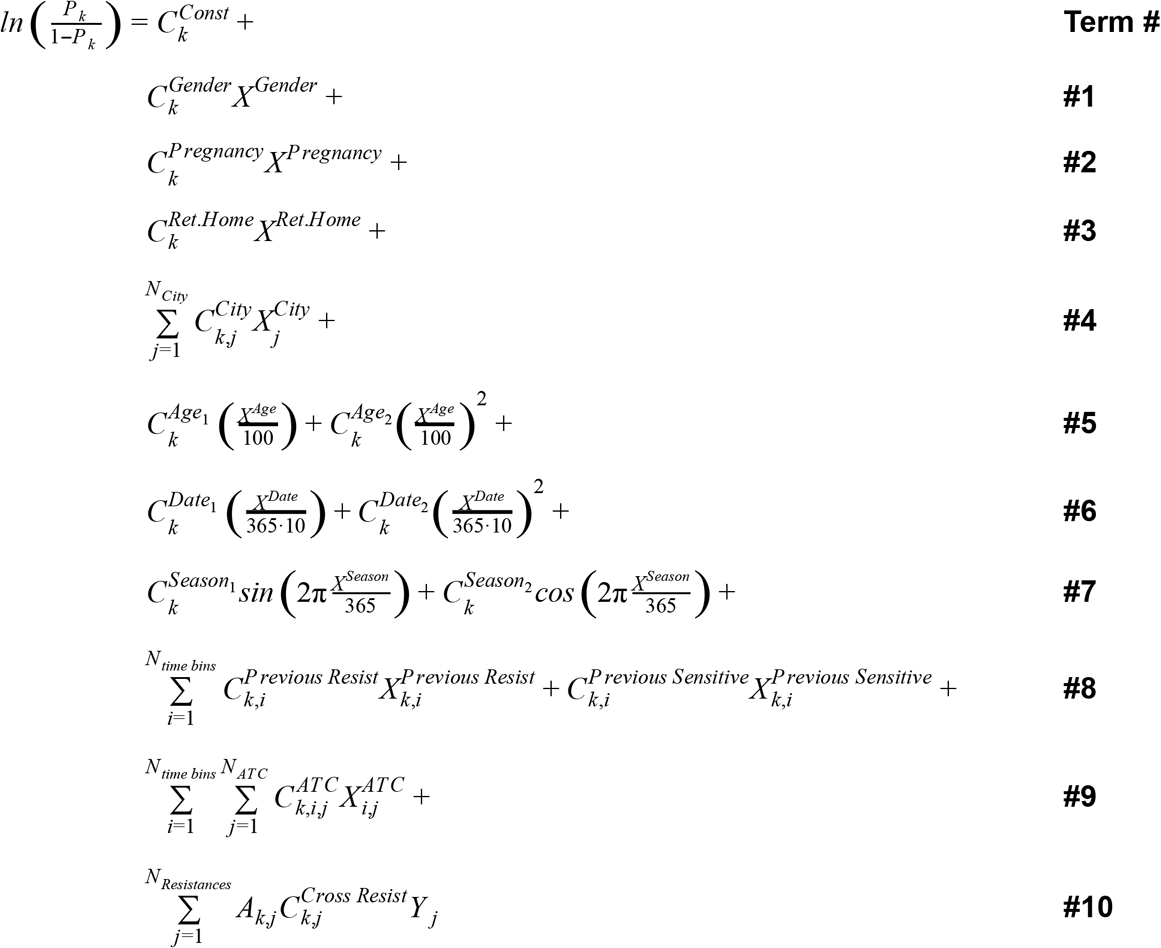

Different combination of the above terms were used in the different regression models as follows (each row in the Table represents a logistic model that was applied to each of the 16 antibiotics):

| **Display item** | **Model name** | **Regression terms** | **Fit data** |
| --- | --- | --- | --- |
| Fig. 2b<br>Sup. Table 1 | Demographics | #1-#7 | All data |
| Sup. Table 1 | Gender, unadjusted | #1 | All data |
| Sup. Table 1 | Pregnancy, adjusted for gender | #2 and #1 | All data |
| Sup. Table 1 | Ret.Home, unadjusted | #3 | All data |
| Sup. Table 1 | City, unadjusted | #4 | All data |
| Sup. Table 1 | Age, unadjusted | #5 | All data |
| Sup. Table 1 | Date, unadjusted | #6 | All data |
| Sup. Table 1 | Season, unadjusted | #7 | All data |
| Fig. 4a,c,d<br>Sup Fig. 6a | Purchase history | #9 | All data |
| Sup Fig. 6b | Purchase history,<br>adjusted for demographics | #9 and #1-#7 | All data |
| Fig. 4b,c,d<br>Sup Fig. 6c | Purchase history,<br>adjusted for cross-resistance | #9 and #10 | All data |
| Fig. 5 | Demographics | #1-#7 | Training range* |
| Fig. 5 | Sample history | #8 | Training range* |
| Fig. 5 | Purchase history | #9 | Training range* |
| Fig. 5 | Complete | #1-#9 | Training range* |

\* See Supplementary Fig. 1 for training range of each of the 16 resistances.

### Calculating odds ratios from logistic regression

For each antibiotic *k*, odds ratios were calculated from the coefficients of above logistic regressions.

#### Binary variables

For the binary variables Gender, Pregnancy and Retirement Home, odds ratio were defined as: 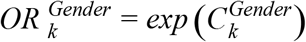 female versus male,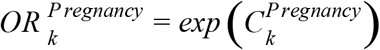 pregnant versus non-pregnant,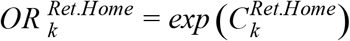 retirement home residence versus patients not residing in retirement homes.

#### Categorical variables

For City, which is a categorical variable, odds ratios for each city *j* relative to the reference (Tel-Aviv) is given by 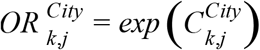, where 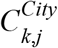 are reported in Supplementary Table 1. In Fig. 2, we report for City the odds ratio 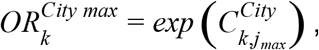, where *j_max_* is the index of the city with largest effect, accounting for error (*j_max_* is defined for each antibiotic *k* as the city *j* with maximal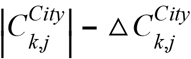, where 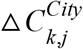 are the standard errors for the regression coefficients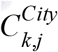

#### Oscillatory variables

For seasonality, the regression coefficients 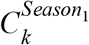 and 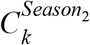 and their CIs are given Supplementary Table 1. In Fig. 2 we provide the odds ratios of the oscillation amplitude: 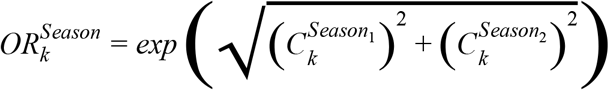.

#### Quadratic variables

For the quadratic variables Age and Date, we again report all the individual regression coefficients and their CIs in Supplementary Table 1. In Fig. 2b, we report, for each antibiotic *k*, effective odds ratios defined as the ratios between the maximal and minimal expected odds taken across the relevant range of these continuous variable (0 ≤ *X^Age^* ≤ 100 and 0 ≤ *X^Date^* ≤ 10 · 365):

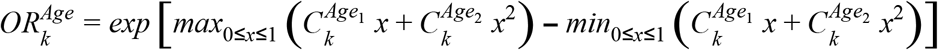

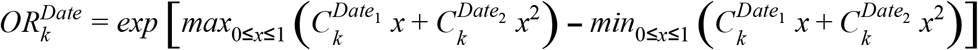

Note that when these quadratic dependencies are monotonic within the relevant range (0 ≤ *x* ≤ 1), the above formulas become simply: 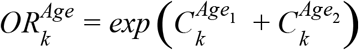,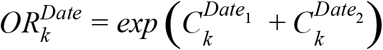.

### Analysis of “memory” across sample pairs

To analyze “memory” of resistance across samples, we considered all pairs of samples from the same patient (across all patients with 2-10 samples) and binned them according to their time difference *t* = *t*_1_ – *t*_2_ (where *t*_1_ and *t*_2_ are the sample dates of the early and late sample; *t* is always negative, indicating information on current sample from past samples) into time bins as indicated by the bars in Fig. 3. In each time bin and for each antibiotic, we counted *N*_*R*→*R*_, *N*_*R*→*S*_, *N*_*S*→*R*_, and *N*_*S*→*S*_ as the number of urine sample pairs where the early and late samples are Resistant, or Sensitive (for example *N*_*R*→*S*_ is the number of same-patient sample pairs, within the time difference bin, where first sample is Resistant and the second Sensitive to the given focal antibiotic. For any antibiotic, only samples for which resistance to the antibiotic was measured were considered). We then calculated for each time difference bin the resistance odds ratio: *OR_pairs_* = (*N*_*R*→*R*_/*N*_*R*→*S*_) / (*N*_*S*→*R*_/*N*_*S*→*S*_).

### Gradient Boosting Decision Trees (GBDT)

GDBT is an ensemble method combining regression trees with weak individual predictive performances, into a single high-performance model. This is done by iteratively fitting decision trees, each iteration targeting the prediction residuals of the preceding tree. The final model is built by combining weighted individual tree contributions, with weights proportional to their performances. For each of the 16 antibiotics, a boosted decision tree ensemble was fitted using all features as defined above (demographics, sample history and drug purchase history) on the training set as defined by the training time period (Supplementary Fig. 1, green bars). This training dataset was sampled to balance resistant/sensitive label frequency. For parameter tuning, a validation dataset was sampled from the training set to be used for model selection (20%). For the estimator of the *i^th^* iteration, a decreasing learning rate η_*i*_ was used such that η_i_ = η_0_α^i^, with an annealing rate α =0.99 and an initial learning rate η_0_ = 0.1. To further promote a diverse ensemble of individual estimators, a 0.9 feature-sampling and observation-sampling rates were used. Fitting of interaction effects is controlled by varying the size of the individual regression trees, with tree estimator of depth *k* producing models with up to k-way interactions. The model was tuned to match data complexity by iteratively increasing tree depth limit of all ensemble estimators while evaluating performance on the validation set, selecting the best depth for each antibiotic.

### Ethical approval

The study protocol was approved by the ethics committee of Assuta Medical Center.

## Contributions

V.S. and R.Kishony perceived the study; I.Y, O.S., G.K., V.S. and R.Kishony designed the study; R.Katz, M.P. and V.S. retrieved and interpreted electronic health records; I. Y., O.S., G.N. and R.Kishony analysed the data; I.Y, O. S., G.C., V.S. and R.Kishony interpreted the results; I.Y and R.Kishony wrote the manuscript with comments from all authors.

## Acknowledgements

We thank M. Datta, A. McAdam, G. Priebe and P. Ramesh for thorough reading of the manuscript and important comments. This work was supported in part by US National Institutes of Health grant R01 GM081617 (to RK) and European Research Council FP7 ERC Grant 281891 (to RK).

---

**Supplementary Figure 1:**
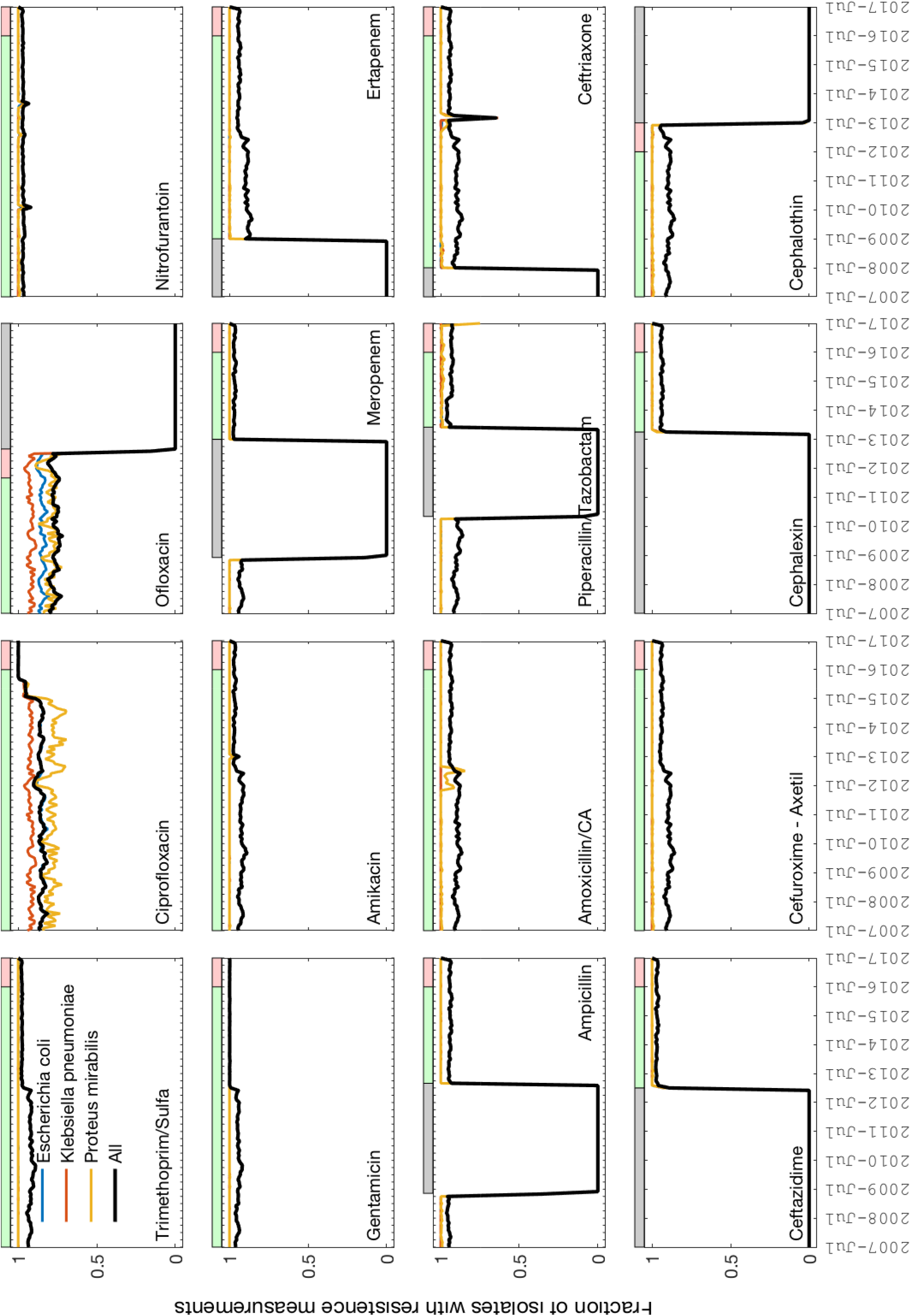
Availability of resistance measurements over time. For each of the 16 antibiotics, the fraction of cultures for which resistance was measured, overall (black) and for each of the three most common species (colors) is plotted across the 10-year sampling period. Availability of measurements determined the time range used for model Training (green horizontal bars) and Testing (red bars) of each resistance. Time periods during which measurements of resistance to a given antibiotic were scarce were removed from analysis (gray bars).

**Supplementary Figure 2:**
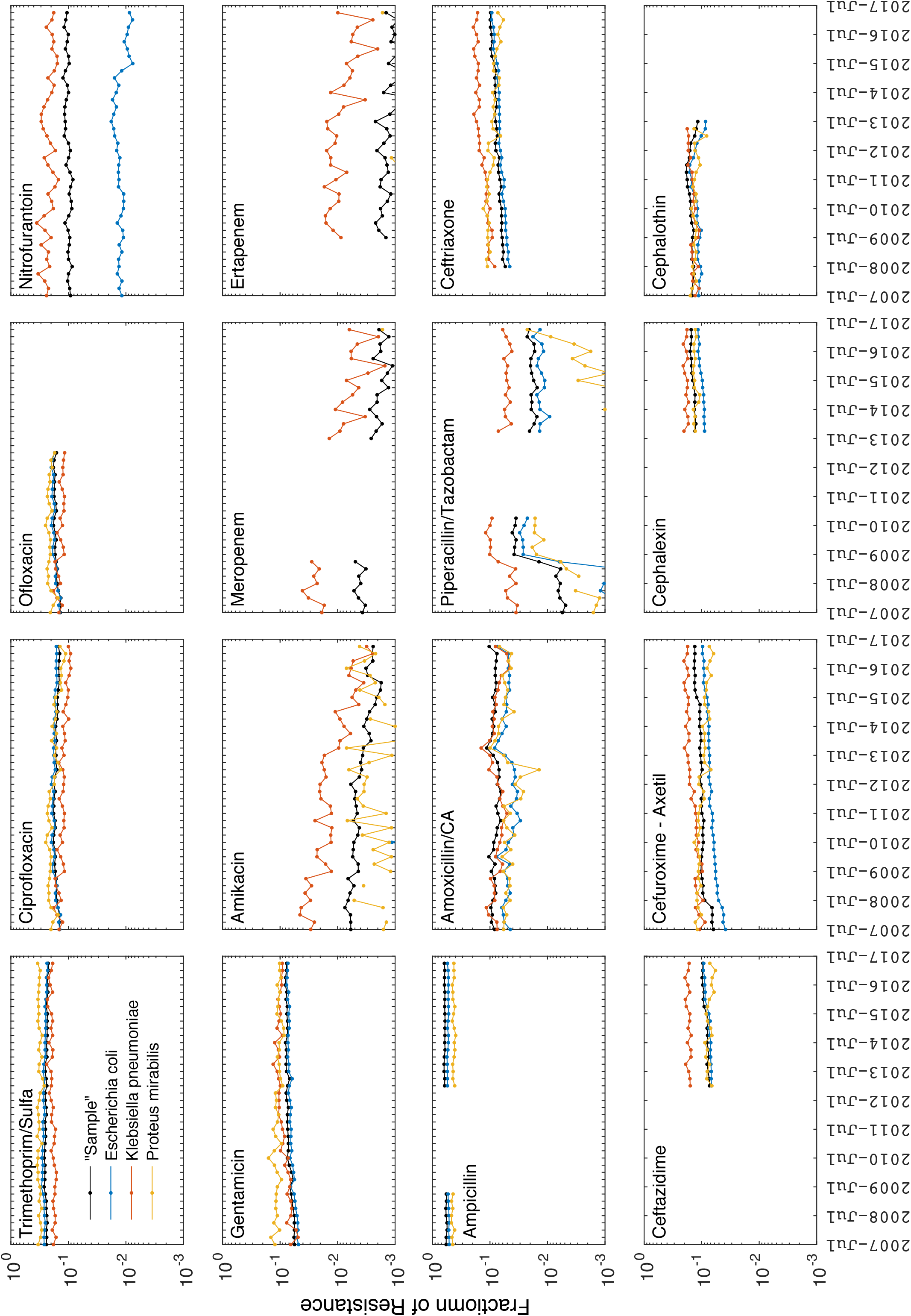
Frequency of resistance over time. Frequencies of resistance of each of the three common species (colored lines) and the overall sample (black lines) over the 10 year sampling period. Empty time intervals correspond to periods during which resistance was not frequently measured (matching the gray horizontal bars of Supplementary Fig. 1). The apparent surge in piperacillin/tazobactam resistance in 2009 is attributed to a switch in VITEK2 reference and does not represent a change in resistance frequency ^43^

**Supplementary Figure 3:**
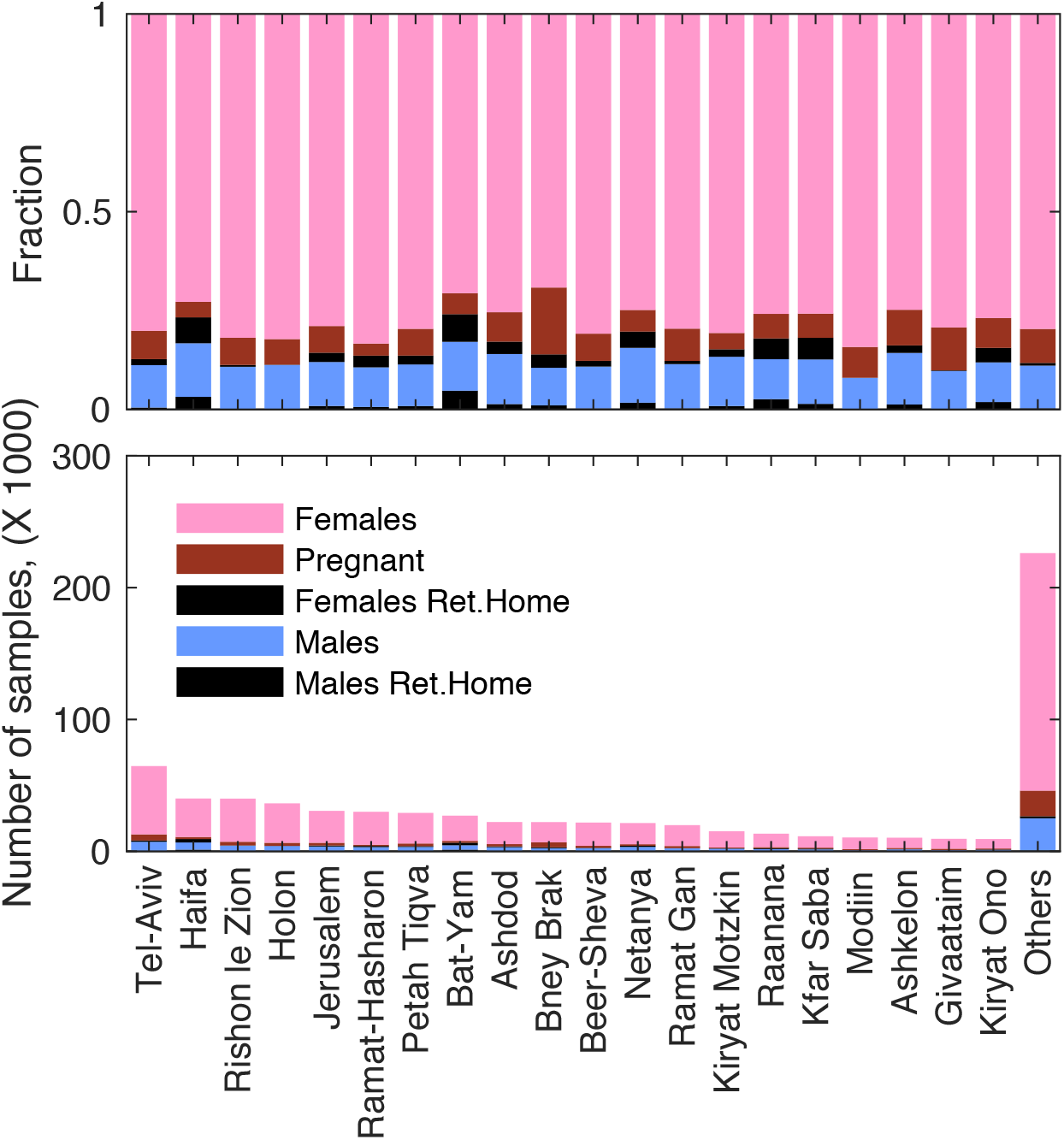
Distribution of UTI demographics by city. Number (bottom) and fraction (top) of UTI samples by 5 demographic categories (legend), for the 20 cities with most samples. ‘Others’ bar for samples in all other cities.

**Supplementary Figure 4:**
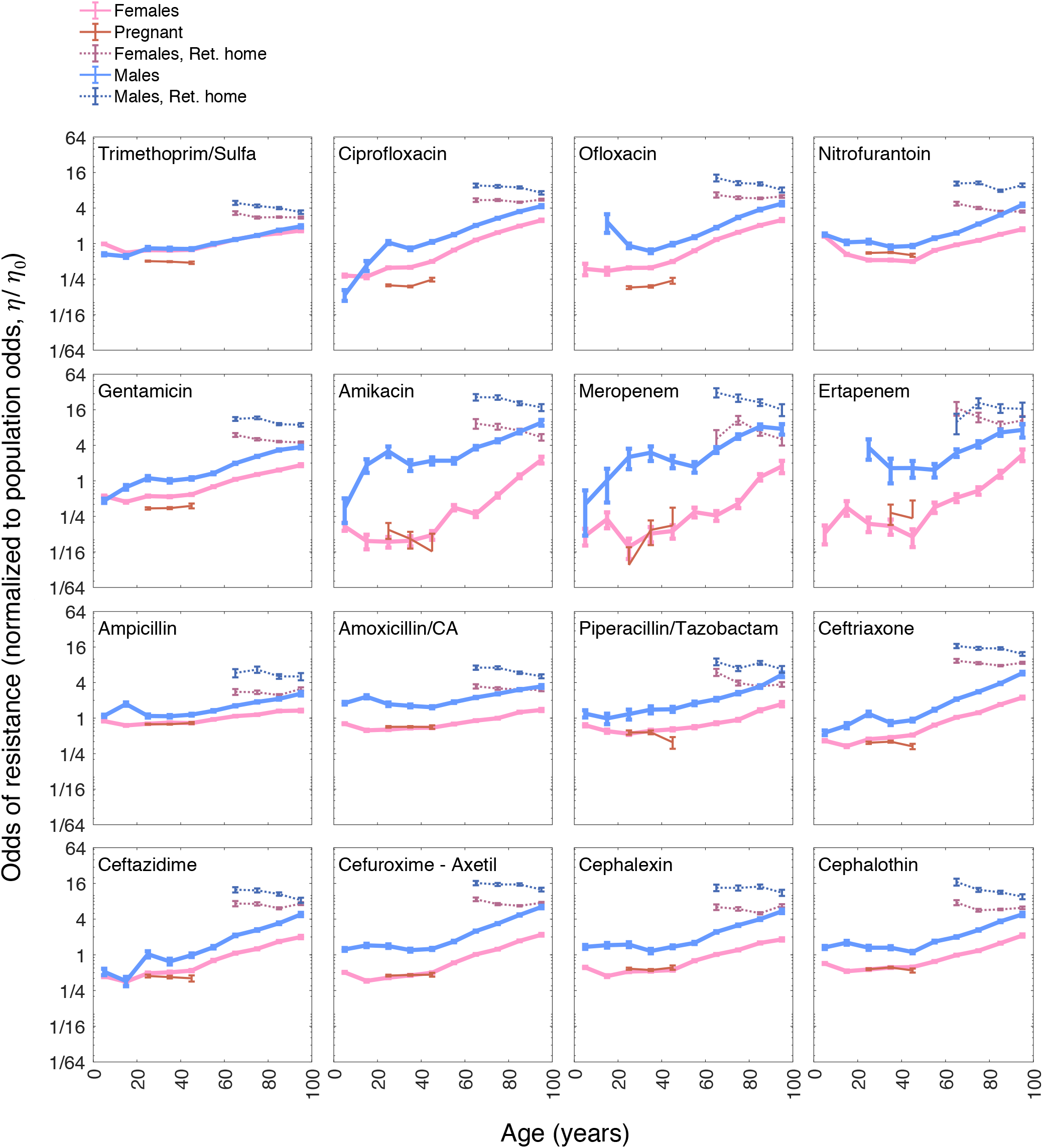
Odds of resistance as a function of age for different demographic groups. Odds of resistance to each of the 16 antibiotics, normalized to the overall population odds (η/η_0_), in each of 10 age bins (0,10,…,100 years) for five non-overlapping demographic groups: men not residing in retirement homes (blue), men residing in retirement homes (dotted blue), women not pregnant and not residing in retirement homes (magenta), women in retirement homes (magenta dotted), and pregnant women (red).

**Supplementary Figure 5:**
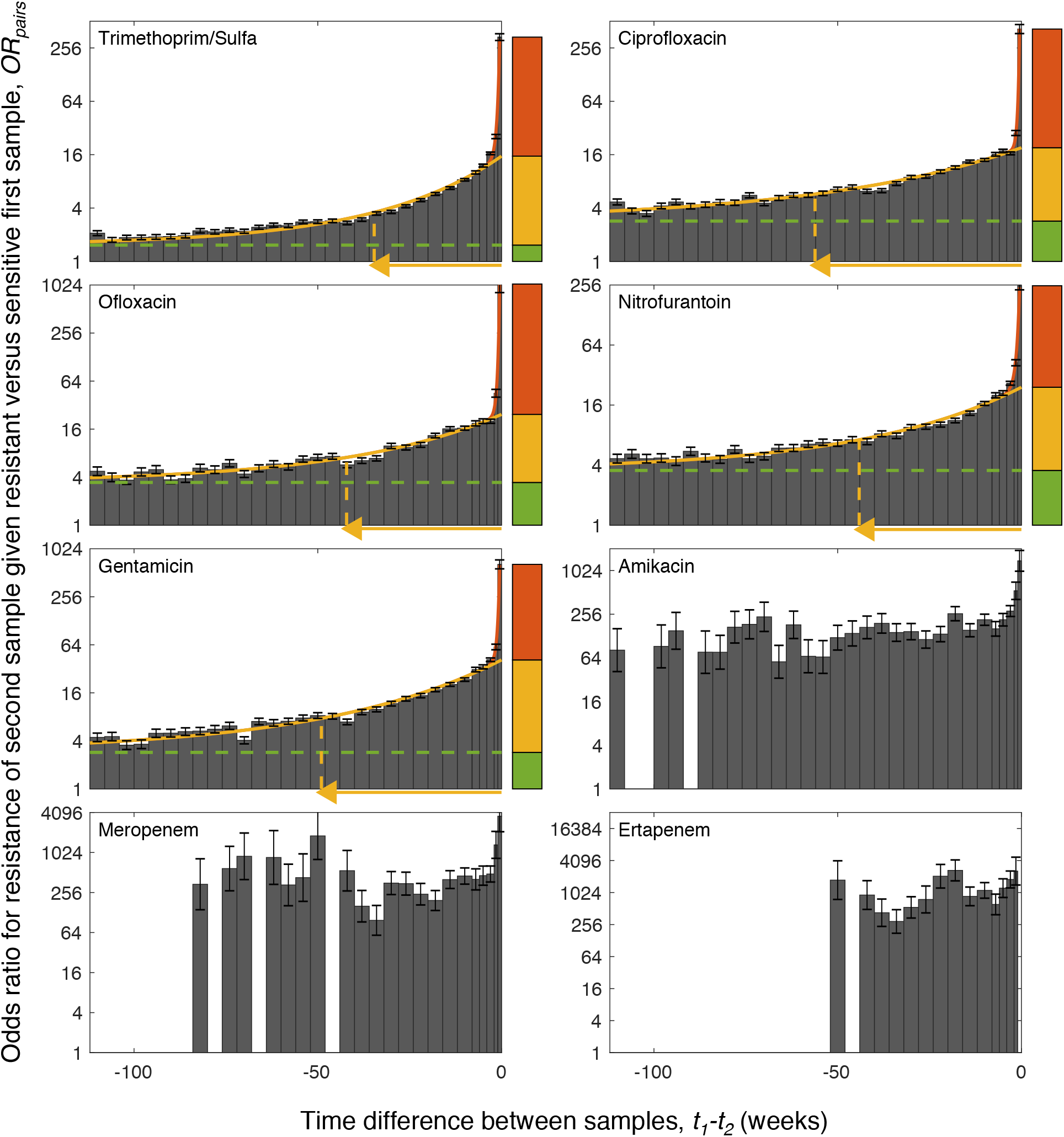
Odds ratio of resistance across same-patient sample pairs. Odds ratio for resistance to each of the 16 antibiotics, given previous resistant versus sensitive urine sample from the same patient, shown as a function of the time between the two samples (*OR_pairs_* = (*N*_*R*→*R*_/*N*_*R*→*S*_) / (*N*_*S*→*R*_/*N*_*S*→*S*_), Methods). Odds ratios are fitted with *OR_pairs_* = *C*_0_*e*^*t*/τ_0_^ + *C*_0_*e*^*t*/τ_m_^ + *C*_∞_, representing short and long memory time scales and a timeless constant (red, yellow and green respectively). The magnitudes of these terms are shown as stacked red, yellow and green bars and the half life (τ_*m*_) of long time scale indicated (yellow arrow). The data of antibiotics amikacin, meropenem, ertapenem and piperacillin/tazobactam were too noisy to be reliably fitted.

**Supplementary Figure 6:**
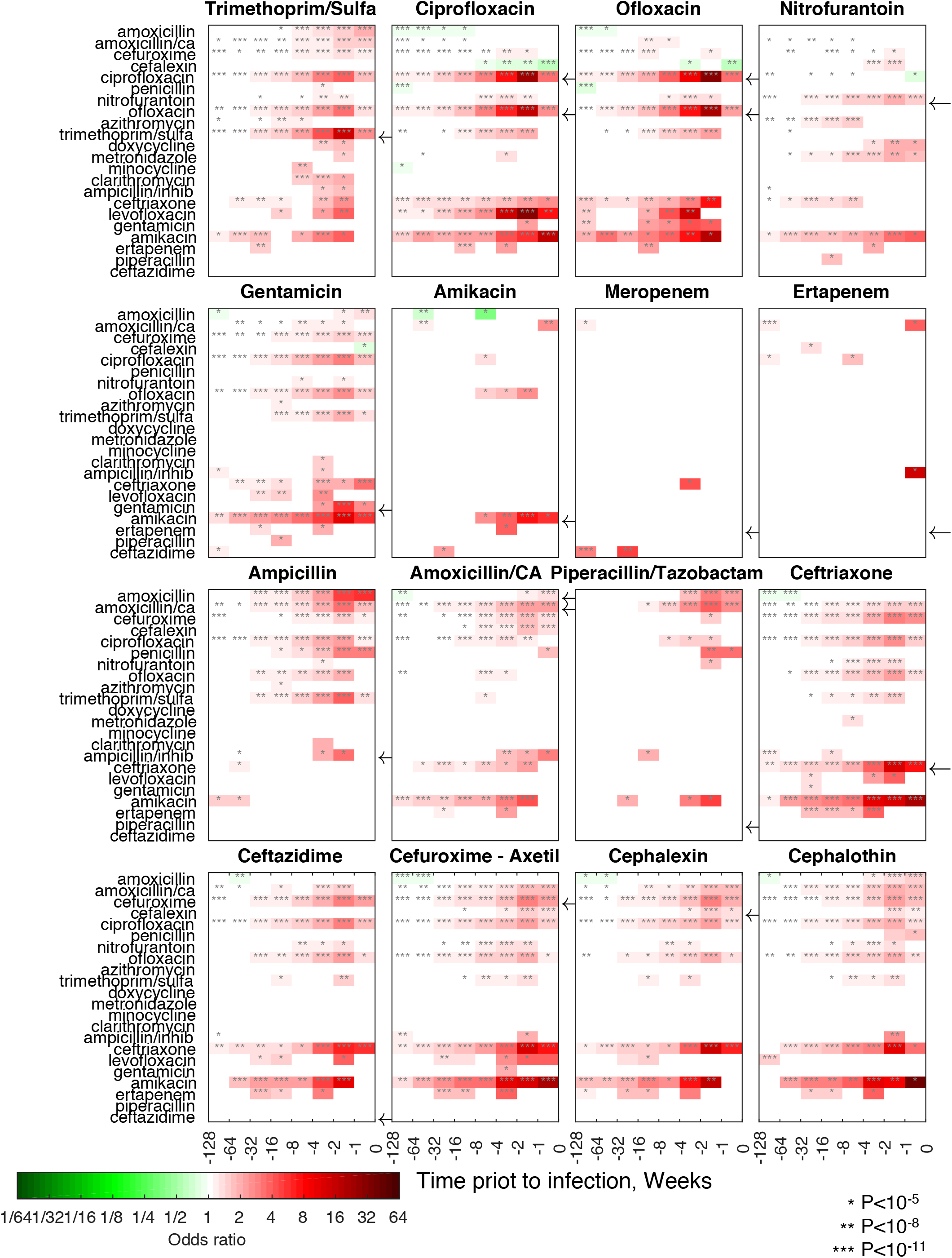
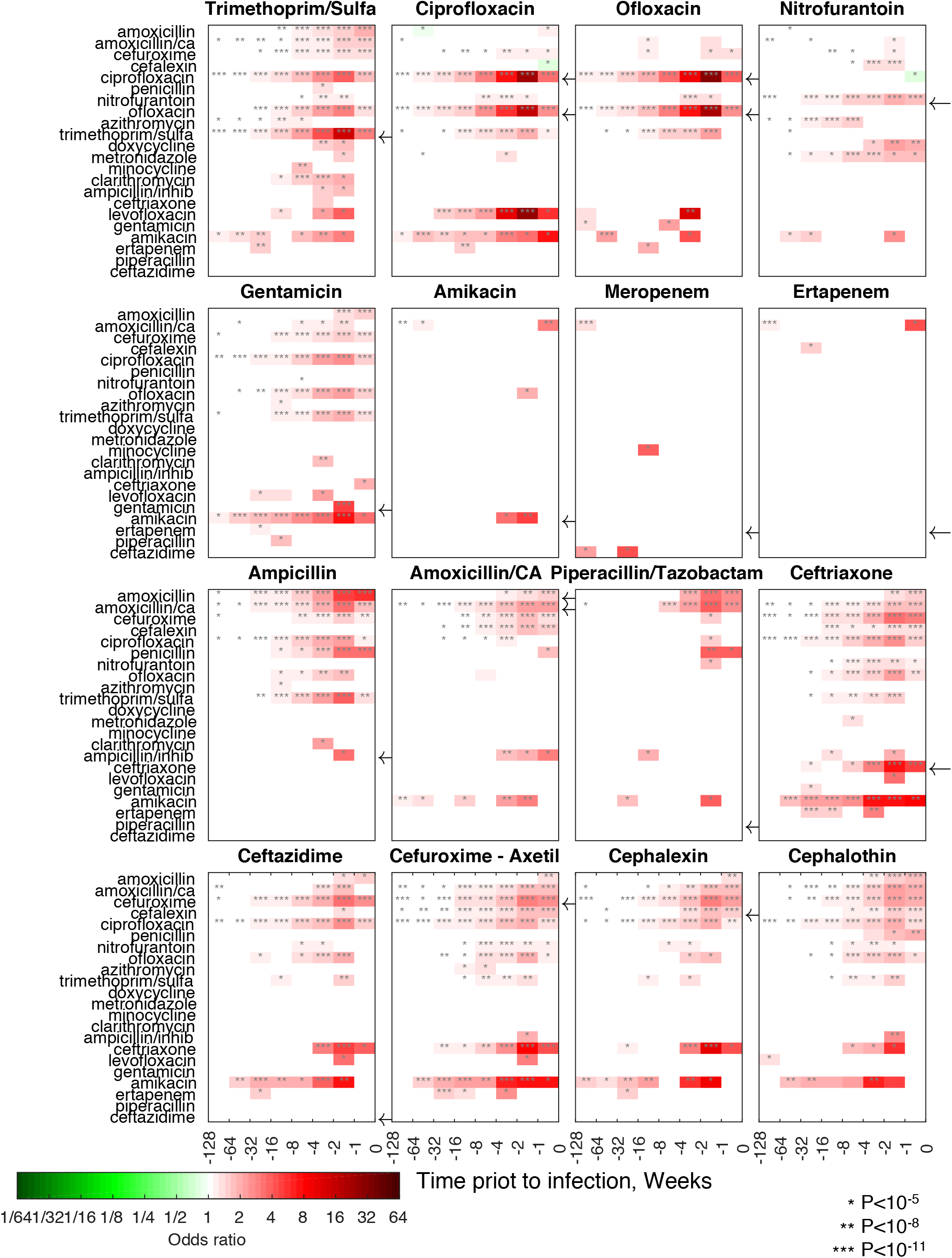
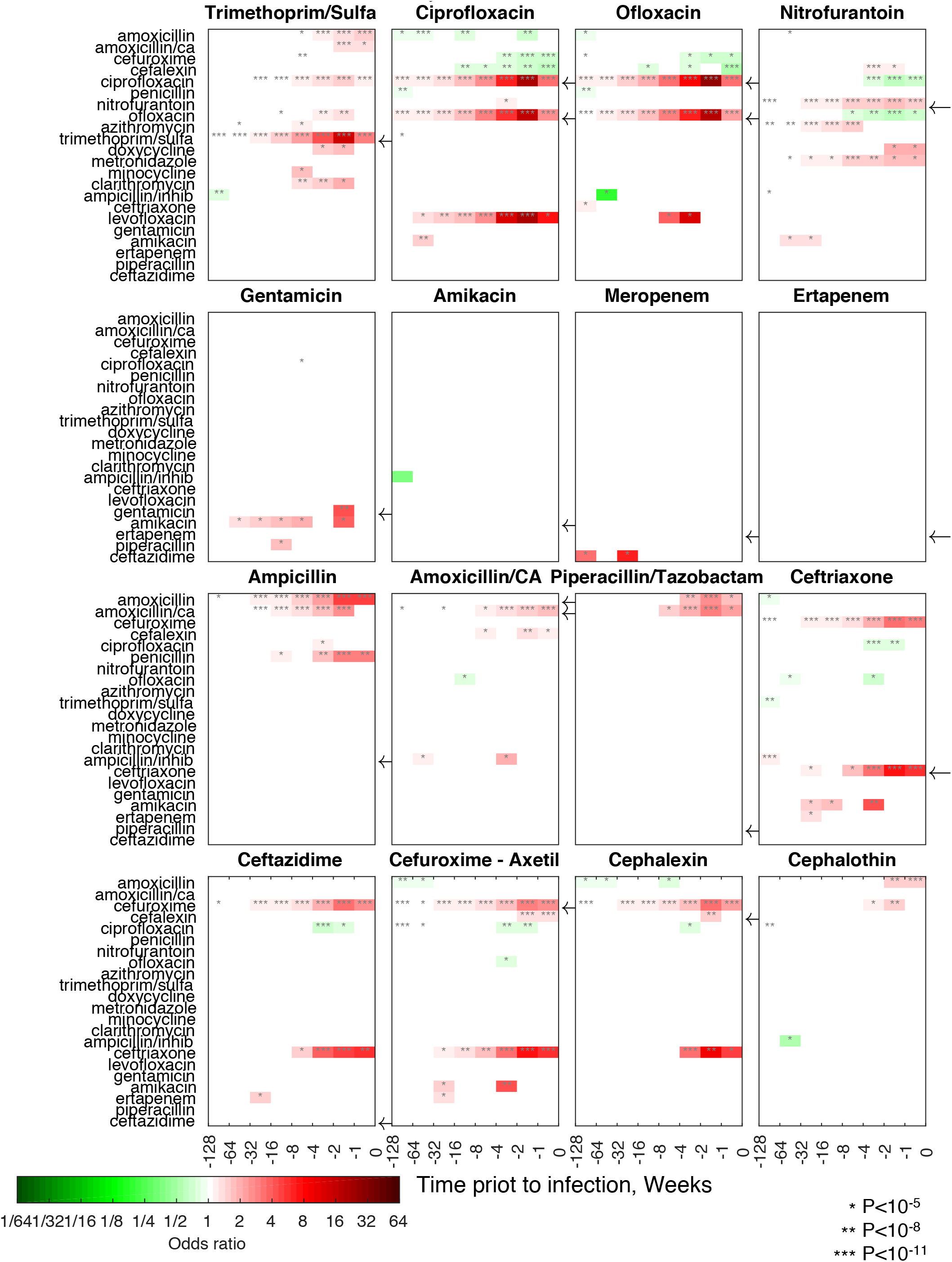
Odds ratios of resistance to each of the antibiotics for past purchases of different drugs across a range of purchase-to-sample time intervals. Adjusted odds ratio of resistance for purchase of antibiotics, 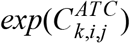 for (**a**) model with purchases only (Methods: Logistic regression “Purchase history”), (**b**) model with purchases and demographics (Methods: Logistic regression “Purchase history adjusted for demographics”), (**c**) model disentangling indirect effects by accounting for linkage between resistances (Methods: Logistic regression “Purchase history adjusted for cross-resistance”). Same graphical scheme as Fig. 4a,b. Gray asterisks indicate statistical significance.

**Supplementary Figure 7:**
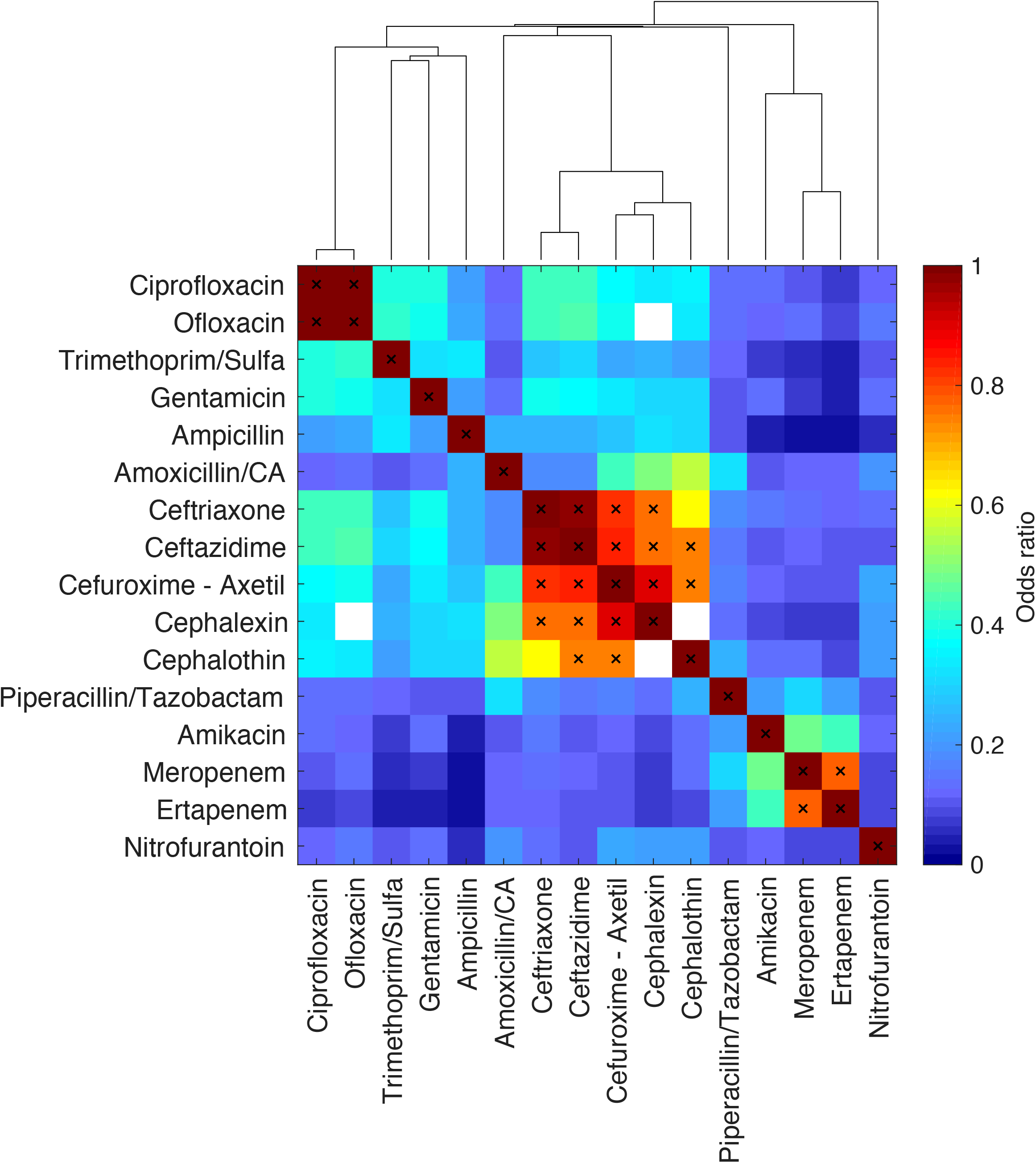
Correlations among resistances to different antibiotics. Correlation among resistance measurements for each pair of antibiotics across all samples for which both resistances were measured (colormap; empty white squares correspond to antibiotic pairs measured in non-overlapping time periods, Supplementary Fig. 1). Correlations values vary between 0.99 (Ciprofloxacin-Ofloxacin) and 0.03 (Ertapenem-Ampicillin). Even the weaker correlations are highly significant (all correlations have P<10^-20^). Pairs of antibiotics with correlation values higher than 0.7 are treated as “analogous”, and their correlations are excluded from analysis of indirect effects of purchases on resistance (Methods: Logistic regression “Purchase history adjusted for cross-resistance”).

**Supplementary Figure 8:**
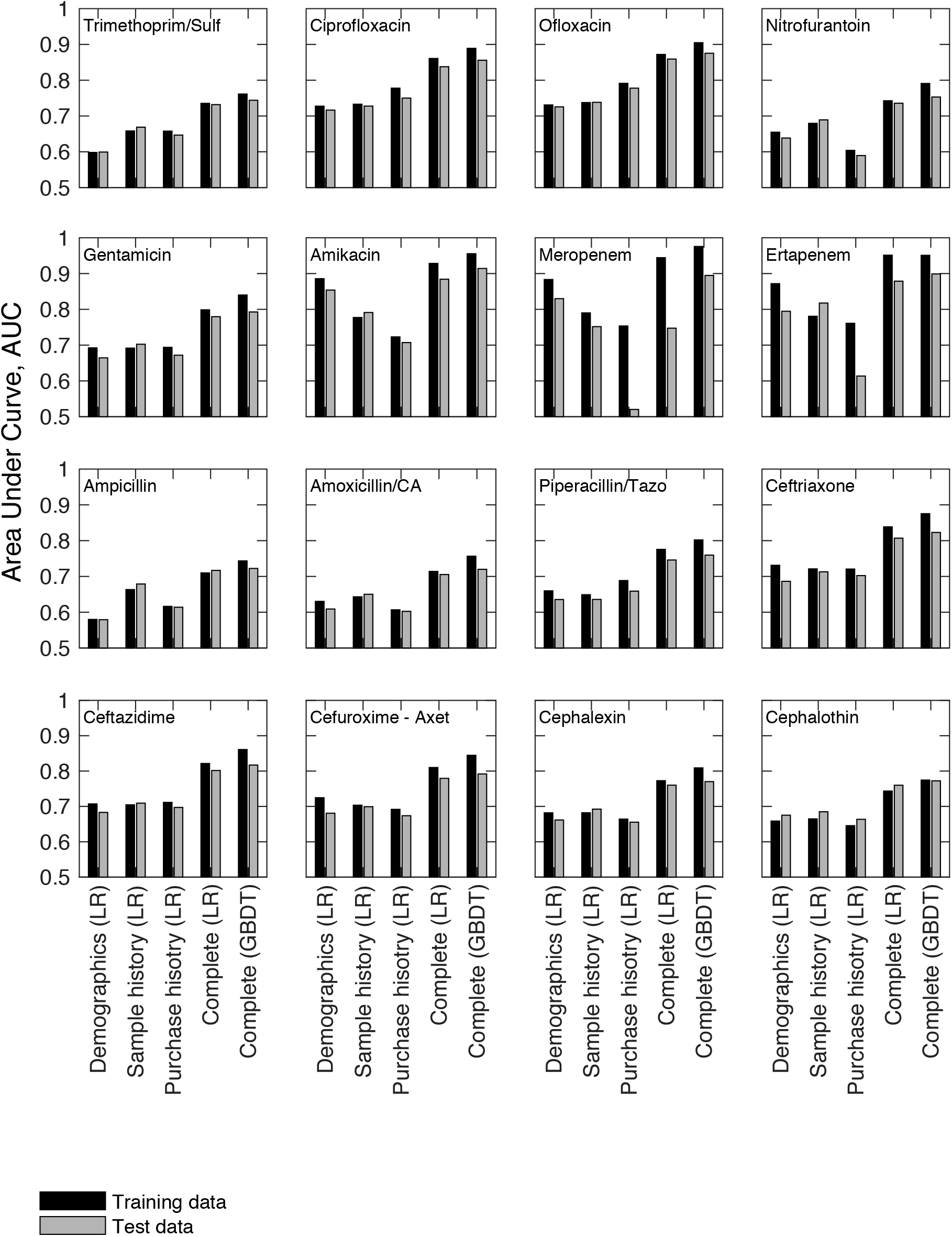
Model performance on test and training data. Area Under Curve (AUC) for Receiver Operator Characteristic for prediction of resistance based on demographics, sample history and purchase history, individually and in a complete model combining all feature sets. Each feature set was modelled using Logistic Regression (LR), and the complete model was modelled by both LR and Gradient Boosting Decision Trees (GBDT). To identify overfitting, model performance on the testing dataset (grey) was contrasted with model performance on the training dataset (black; Supplementary Fig. 1 for definition of training and test time periods). For most antibiotics both LR and GBDT models had similar predictive power when tested on the training and the testing datasets, suggesting no significant overfitting. We could however detect overfitting for the LR model of meropenem, especially in the purchase history model, corresponding to the small number of purchases of this drug (Supplementary Table 2). This overfitting was substantially reduced in the GBDT model, where the hyperparameter tuning determined a relatively low optimal decision tree depth limit (3 compared to 6-7 for most other antibiotics), lowering the variance error, and reducing overfitting.

**Supplementary Figure 9:**
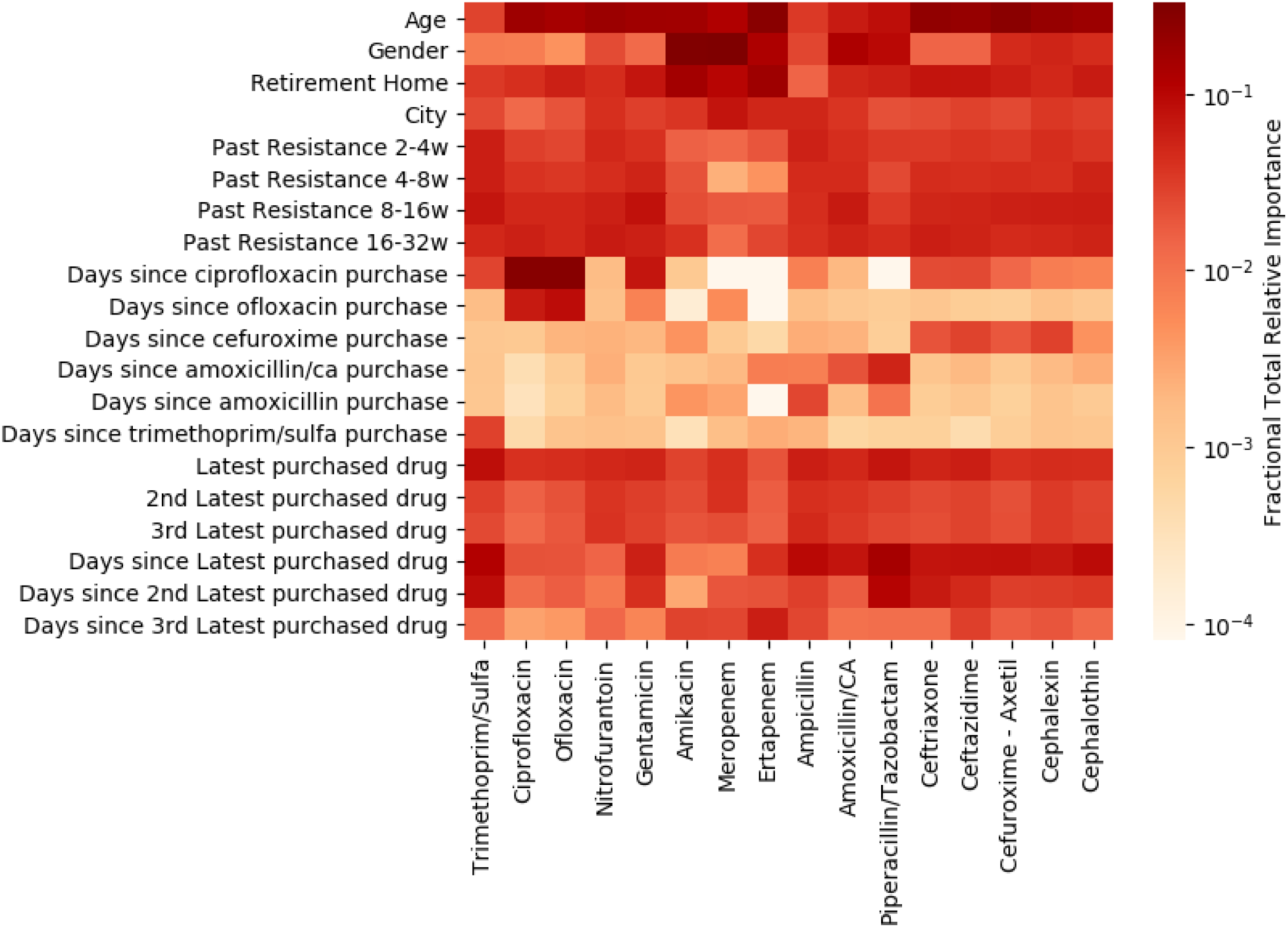
Feature importance in Gradient Boosting Decision Trees (GBDT) The relative importance of each feature on the prediction of the GBDT model of each antibiotic is shown (color map) ^44^ Importance is defined, for each feature, as its fractional estimated total error improvement, which is the sum of error difference over all instances of the feature in all trees in an ensemble ^44^ For each antibiotic, total feature error improvements are normalized to one. The features shown are those of highest influence for each of the main features sets: demographics, sample history and drug purchase history.

**Supplementary Table 1:**
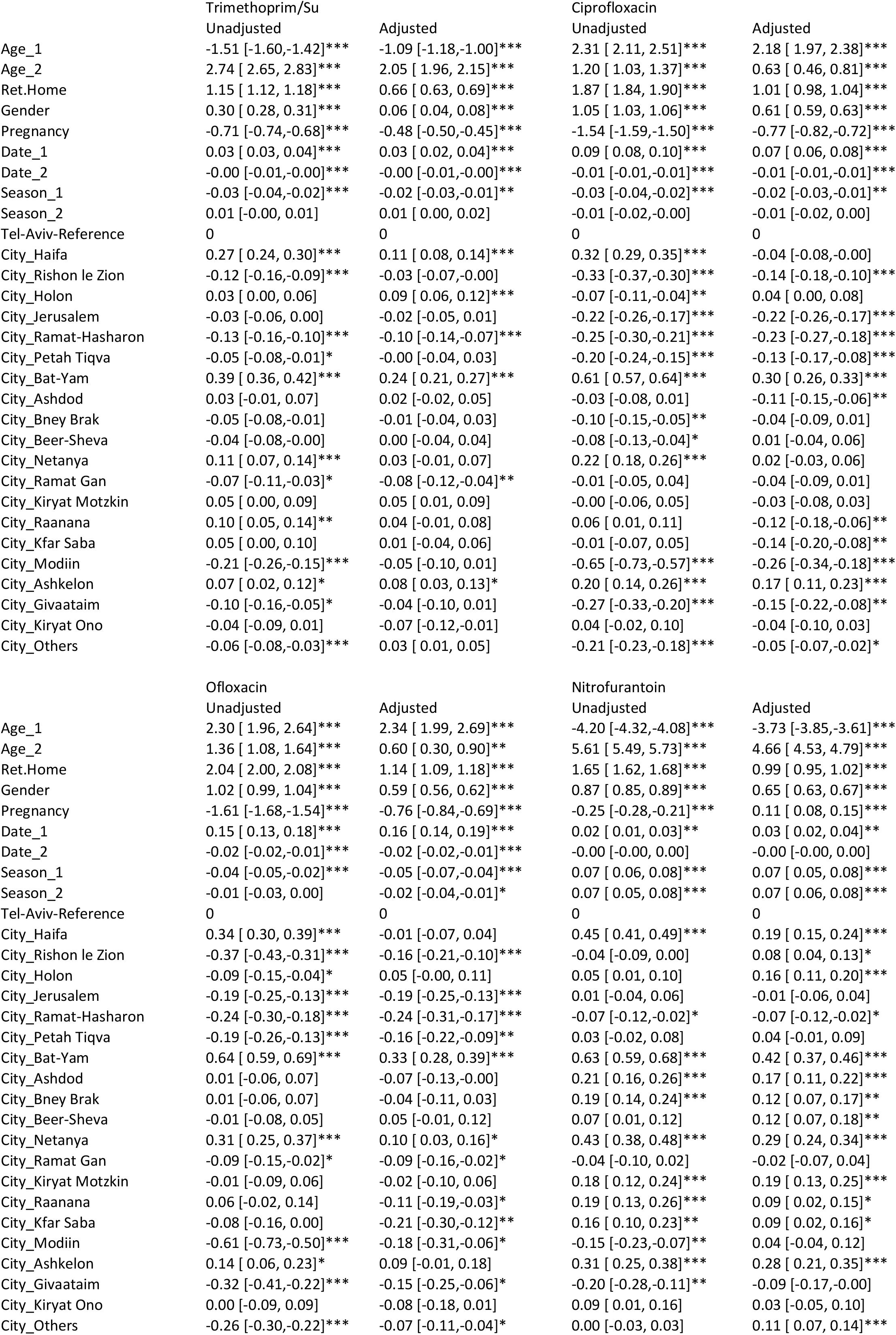
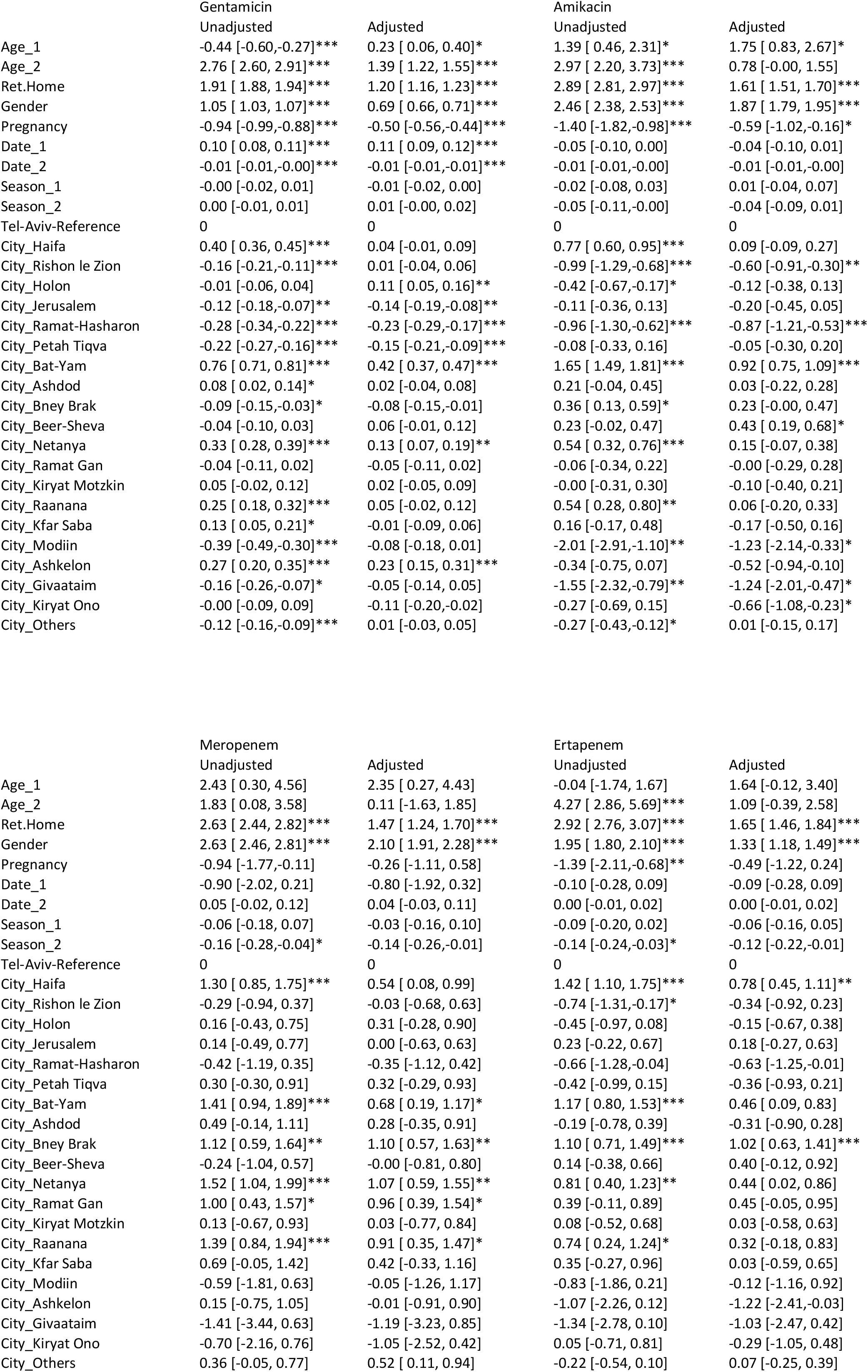
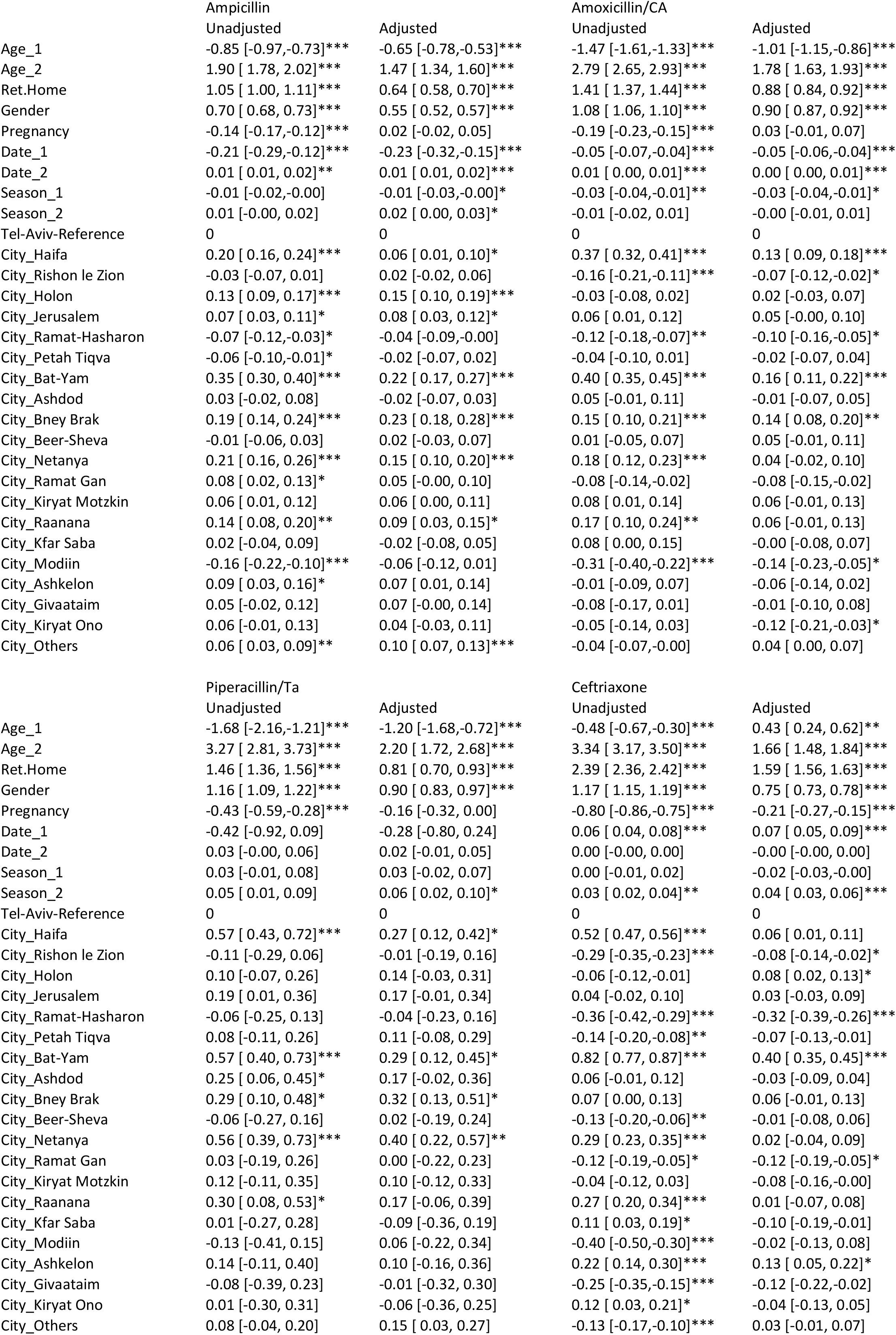
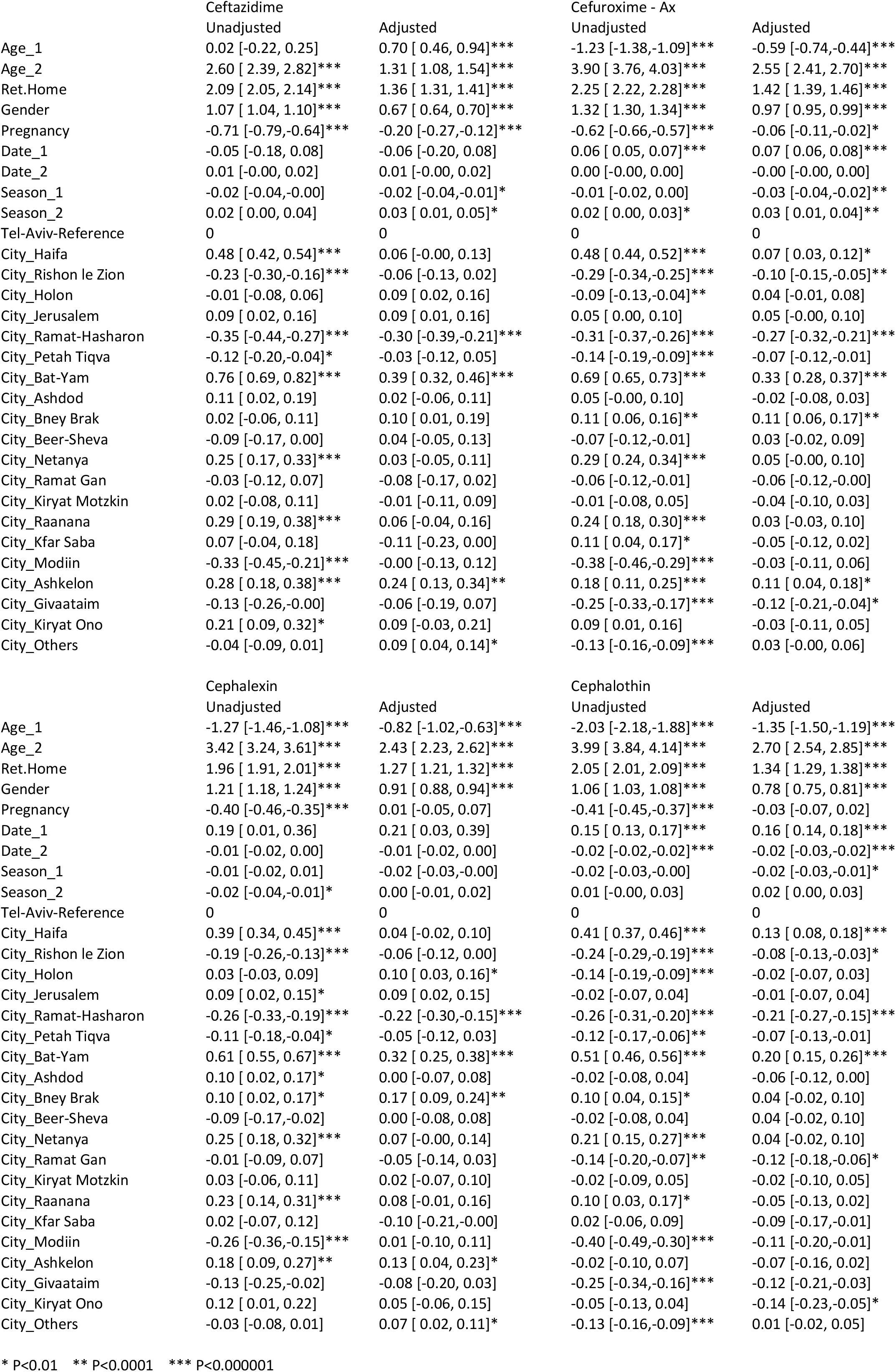
Regression coefficients for association of resistance with demographics. Logistic regression of resistance to each of the 16 antibiotics with respect to each of the demographic features alone (unadjusted) and when accounted for together (adjusted). For definitions of listed demographic terms see Methods.

**Supplementary Table 2:** Antimicrobial drugs considered in purchase model.

| Group | Name | ATC codes | Counts |
| --- | --- | --- | --- |
| 1 | amoxicillin | J01CA04 | 1321804 |
| 2 | amoxicillin/ca | J01CR02 | 743798 |
| 3 | cefuroxime | J01DC02 | 552986 |
| 4 | cefalexin | J01DB01 | 507924 |
| 5 | ciprofloxacin | J01MA02 | 458447 |
| 6 | penicillin | J01CE01 J01CE02 J01CE08 | 363346 |
| 7 | nitrofurantoin | J01XE01 | 321099 |
| 8 | roxithromycin | J01FA06 | 287591 |
| 9 | ofloxacin | J01MA01 | 265931 |
| 10 | azithromycin | J01FA10 | 233245 |
| 11 | trimethoprim/sulfa | J01EE01 | 232708 |
| 12 | fosfomycin | J01XX01 | 175813 |
| 13 | doxycycline | J01AA02 | 172187 |
| 14 | metronidazole | J01XD01 | 116291 |
| 15 | minocycline | J01AA08 | 94004 |
| 16 | clarithromycin | J01FA09 | 68021 |
| 17 | clindamycin | J01FF01 | 52890 |
| 18 | erythromycin | J01FA01 | 48139 |
| 19 | methenamine | J01XX05 | 39459 |
| 20 | ampicillin/inhib | J01CA51 J01CR01 | 33696 |
| 21 | cefaclor | J01DC04 | 22971 |
| 22 | cloxacillin | J01CF02 | 21678 |
| 23 | ceftriaxone | J01DD04 | 14317 |
| 24 | cefixime | J01DD08 | 13944 |
| 25 | levofloxacin | J01MA12 | 13944 |
| 26 | tetracycline | J01AA07 | 7906 |
| 27 | ampicillin | J01CA01 | 4793 |
| 28 | fusidic acid | J01XC01 | 4265 |
| 29 | gentamicin | J01GB03 | 3087 |
| 30 | vancomycin | J01XA01 | 2053 |
| 31 | amikacin | J01GB06 | 2000 |
| 32 | ertapenem | J01DH03 | 1703 |
|  | rifaximin | A07AA11 | 1645 |
| 11 | sulfamethoxazol | J01EC01 | 1567 |
|  | cefazolin | J01DB04 | 1171 |
|  | colistin | J01XB01 | 1156 |
| 34 | ceftazidime | J01DD02 | 919 |
| 32 | meropenem | J01DH02 | 849 |
| 33 | piperacillin/inhib | J01CR05 | 725 |
|  | neomycin | A07AA01 | 704 |
|  | spiramycin | J01FA02 | 643 |
|  | clofazimine | J04BA01 | 630 |
|  | nalidixic acid | J01MB02 | 568 |
| 32 | imipenem/inhib | J01DH51 | 412 |
|  | norfloxacin | J01MA06 | 377 |
|  | dapsone | J04BA02 | 224 |
|  | moxifloxacin | J01MA14 | 197 |
|  | tobramycin | J01GB01 | 142 |
| 33 | piperacillin | J01CA12 | 127 |

